# Dynamic changes in the urine proteome of tumor-bearing mouse models of B16 melanoma and RM-1 prostate cancer

**DOI:** 10.1101/2020.04.03.023366

**Authors:** Lujun Li, Xuanzhen Pan, Yongtao Liu, Ting Wang, Youhe Gao

## Abstract

Urine can accumulate changes and reflect early physiological and pathological changes of various diseases, such as tumors. Therefore, urine is an ideal source for identification of early biomarkers. In this study, melanoma and prostate cancer-bearing mouse models were established by subcutaneous injection of B16 and RM-1 cells, respectively. Urine samples were collected at four time points during tumor growth. Based on data-independent acquisition (DIA) technology, liquid chromatography-tandem mass spectrometry (LC-MS/MS) was used for quantitative analysis. Compared with those before the injection of B16 cells, 38 human homologous differential proteins were identified, and 18 proteins were reported to be related to melanoma. Before the tumor was visible, there were 4 differential proteins, and all were reported to be related to melanoma. Compared with that before the injection of RM-1 cells, a total of 14 human homologous differential proteins were identified, and 9 proteins were reported to be associated with prostate cancer. Before the tumor was palpable, 9 proteins showed significant differences. There were significant differences between the two tumor-bearing models. Through the above experiments and analysis, we found that the urine proteome can reflect the changes in the development and provide early biomarkers of the two tumors and provide clues for the early clinical diagnosis of these diseases.

---

Skin melanoma is one of the most common cancers in the world[1]. Melanoma is produced by the malignant transformation of melanocytes. This disease can occur in any part of the skin and mucous membranes. Melanoma easily metastasizes and is highly invasive and malignant, leading to high mortality. The diagnosis of melanoma is based on a combination of clinical and pathological techniques, including physical examination, histopathological examination and imaging examination. However, due to its variable histological morphology, with little or even no visible pigment, it is difficult to diagnose patients in the early stage and distinguish this condition from benign nevus in the clinic, which makes its early diagnosis difficult[2]. Early treatment of melanoma is mainly surgery, and advanced treatment includes chemotherapy, targeted therapy and immunotherapy. Compared with chemotherapy alone, targeted therapy and immunotherapy have significantly improved the survival rate of patients with advanced melanoma, but these two therapies are usually limited by the drug resistance rate and are not universally applicable[3, 4]. The absolute survival rate of patients is still very low[5]. The 5-year survival rates of skin melanoma at stage I to IV are 97% (stage IA), 84% (stage IB), 68%, 55%, and 17%[6]. Therefore, early diagnosis can lead to early treatment and improve the survival rate of patients.

Prostate cancer is one of the most common malignant tumors in men. In Europe and America, the incidence rate is highest among those of all cancers in males. The mortality rate is second only to that of lung cancer, ranking second for malignant tumors in males[7]. The incidence rate and mortality rate of the Asian population have also shown an increasing trend[8]. Prostate cancer grows slowly, has no clinical symptoms in the early stage, is easily confused with benign hyperplasia, and develops rapidly in the later stage, leading to malignant metastasis[9, 10]. Early detection and treatment can significantly improve the survival rate of prostate cancer[11]. Existing diagnostic methods, such as digital rectal examination (DRE), transrectal ultrasound (TRUS), serum PSA, and puncture biopsy, cannot truly achieve high sensitivity and specificity. Therefore, it is necessary to find a new method for the early diagnosis of prostate cancer.

Biomarkers are a class of indicators that can objectively reflect normal physiological and pathological processes[12]. From a clinical perspective, biomarkers can help with the monitoring, prediction and diagnosis of multifactorial diseases at different stages[13]. Blood is a relatively stable system. As the disease progresses, when the blood reaches the critical point of compensation, loss of compensation is observed. Before reaching the critical point of decompensation, changes caused by the disease are eliminated by the blood. In this process, one of the most important environments for metabolic waste is urine. Therefore, sensitive early biomarkers are more likely found in the urine than the blood[14].

Urine is susceptible to gender, age, diet and other factors[15]. The use of animal models with controllable living conditions and clear genetic backgrounds can not only avoid the interference of the above factors but can also continuously monitor the course of disease. Previous studies have shown that urine can sensitively reflect the changes in tumor cells in animal models[16, 17] To explore the effects of the occurrence and development of melanoma and prostate cancer on the urine proteome, we established tumor-bearing mouse models by subcutaneously injecting B16 and RM-1 tumor cells, respectively. Urine samples were collected at four time points corresponding to tumorigenesis and development, and the urine proteome was analyzed by liquid chromatography-tandem mass spectrometry (LC-MS/MS) based on data-independent acquisition (DIA) technology. The above results provide valuable clues for the early clinical diagnosis of melanoma and prostate cancer in the future.

## 1 Materials and methods

### 1.1 Cell lines and reagents

B16 mouse melanoma cells and RM-1 mouse prostate cancer cells were purchased from the Cell Bank of the Chinese Academy of Sciences. The cells were cultured in RPMI-1640 (Corning) containing 10% fetal bovine serum (Gibco) at 37°C and 5% CO2.

### 1.2 Experimental animals and model establishment

Seven-week-old male C57BL/6 mice were purchased from Beijing Vital River Laboratory Animal Technology Co., Ltd. The animal license is SCXK (Beijing) 2016-0006. Animal procedures were approved by the Institute of Basic Medical Sciences Animal Ethics Committee, Peking Union Medical College (ID: ACUC-A02-2014-008). All animals were housed in a standard environment with a 12-h light-dark cycle under controlled indoor temperature (22 ± 2°C) and humidity (65–70%).

The methods of subcutaneous injection of B16[18–20] and RM-1[21, 22] cells to establish the tumor-bearing mouse model are as follows: the number of living cells was more than 95% as calculated by trypan blue staining. (1) The melanoma-bearing mouse model was established by subcutaneous injection of 0.1 mL of B16 cells (1.8×10^5^) in the right hind limb of the mice (n = 8). (2) The model of prostate cancer-bearing mice was established by subcutaneous injection of 0.1 mL RM-1 cells (1.9×10^4^) into the right hind limb of mice (n = 4). This experiment used an autocontrol, the urine collected before the injection of tumor cells was used as the control group, the time was recorded as day 0, and the time of subcutaneous injection of the tumor cells was recorded as day 1.

### 1.3 Urine collection and sample preparation

Urine samples of the B16 tumor-bearing mice on days 0, 4, 7 and 14 and the RM-1 tumor-bearing mice on days 0, 7, 15 and 22 were collected. The mice were placed in a metabolic cage alone overnight (12 h) to collect urine samples. During urine collection, no food was provided, and free drinking water was available to avoid urine pollution.

The collected urine was centrifuged at 4°C and 3000 × g for 10 min to remove the cells and pellets. The supernatant was stored at −80°C. Prior to urine protein extraction, the urine samples were centrifuged at 4°C and 12000 × g for 10 min to remove cell debris. Four volumes of precooled ethanol were used, and the supernatant was precipitated at 4°C for 12 h. The above samples were centrifuged at 4°C and 12000 × g for 10 min. The pellet was resuspended in lysate (8 mol/L urea, 2 mol/L thiourea, 25 mmol/L DTT and 50 mmol/L Tris). The Bradford method was used to determine the protein concentration.

Filter-aided sample preparation (FASP)[23] was used for membrane-assisted enzymolysis of urine proteins. One hundred micrograms of protein was loaded onto 10-kD cutoff filter devices (Pall, Port Washington, NY). The samples were washed sequentially with UA (8 mol/L urea, 0.1 mol/L Tris-HCl, pH 8.5) and 50 mmol/L NH4HCO3. Then, 20 mmol/L DTT was added to reduce the protein (37°C, 1 h), and 50 mmol/L IAA was reacted in the dark for 30 min to alkylate the proteins. The proteins were digested with trypsin (Trypsin Gold, Promega, Fitchburg, WI, USA) (enzyme-to-protein ratio of 1:50) for 14 h at 37°C. The peptides were collected and desalted on Oasis HLB cartridges (Waters, Milford, MA) and then dried by vacuum evaporation (Thermo Fisher Scientific, Bremen, Germany) and stored at −80°C.

Pooled peptides were fractionated by a high-pH reversed-phase peptide fractionation kit (84868, Thermo Fisher, USA) according to the manufacturer’s instructions. The peptide samples were eluted with a gradient of increasing acetonitrile concentration. A total of 10 fractions were collected: the flow-through fraction, wash fraction and 8 step gradient sample fractions. The fractionated samples were dried completely and resuspended in 20 μL of 0.1% formic acid.

### 1.4 LC-MS/MS analysis

#### 1.4.1 DDA-MS

LC-MS/MS data acquisition was performed on a Fusion Lumos mass spectrometer (Thermo Scientific, Germany) coupled with an EASY-nLC 1200 high-performance liquid chromatography system (Thermo Scientific, Germany). For both DDA-MS and DIA-MS modes, the same LC settings were used for stability of the retention time. The digested peptides were dissolved in 0.1% formic acid and loaded on a trap column (75 µm × 2 cm, 3 µm, C18, 100 Å), and the eluent was transferred to a reversed-phase analytical column (50 µm × 250 mm, 2 µm, C18, 100 Å) with an elution gradient of 5-30% phase B (79.9% acetonitrile, 0.1% formic acid, flow rate of 0.5 μL/min) for 90 min. For fully automated and sensitive signal processing, the calibration kit (iRT kit, Biognosys, Switzerland) was spiked at a concentration of 1:20 v/v in all samples.

For the generation of the spectral library, 10 peptide fractions were analyzed in DDA-MS mode. The parameters were set as follows: the full scan was acquired from 350 to 1 550 m/z at 60 000, the cycle time was set to 3 s (top speed mode), the auto gain control (AGC) was set to 1e6 and the maximum injection time was set to 50 ms. The MS/MS scans were acquired in the Orbitrap at a resolution of 15000 with an isolation window of 2 Da and collision energy at 32% (HCD). The AGC target was set to 5e4, and the maximum injection time was 30 ms.

#### 1.4.2 DIA-MS

For the DIA-MS method, 48 individual samples were analyzed in DIA mode. The parameters were set as follows: the full scan was acquired from 350 to 1 550 m/z at 60000, followed by DIA scans with a resolution of 30000, HCD collision energy of 32%, AGC target of 1e6 and maximal injection time of 50 ms. Thirty-six variable isolation windows were developed, and the calculation method of the windows was as follows: the DDA search results were sorted according to the number of identified peptide segments by m/z and divided into 36 groups. The m/z range of each group is the window width for collecting DIA data.

### 1.5 Data analysis

For generation of the spectral library, the ten fractions’ raw data files acquired by the DDA mode were processed using Proteome Discoverer (version 2.3, Thermo Scientific, Germany) with the SwissProt mouse database (released in May 2019, containing 17016 sequences) appended with the iRT peptide sequences. The search parameters were as follows: parent ion mass tolerance, 10 ppm; fragment ion mass tolerance, 0.02 Da; fixed modifications, carbamidomethylated cysteine (+58.00 Da); and variable modifications, oxidized methionine (+15.995 Da). Other settings included the default parameters. The applied false discovery rate (FDR) cutoff was 0.01 at the protein level. The results were then imported to Spectronaut™ Pulsar (Biognosys, Switzerland) software to generate the spectral library: 798 proteins and 28356 fragments.

The DIA-MS raw files were imported to Spectronaut™ Pulsar with the default settings. Quantitative analysis was based on the peak areas of all fragment ions for MS2. Each protein was identified with at least two specific peptides and a Q value < 0.01. The screening criteria for differential proteins were as follows: fold change of protein abundance > 1.5, p value < 0.01 by t-tests.

### 1.6 Functional enrichment analysis

The online database DAVID 6.8 (https://david.ncifcrf.gov/) was used to perform the functional annotation of the differential proteins, including molecular function, cell component and biological process. Pathway analysis of differential proteins was performed using IPA software (Ingenuity Systems, Mountain View, CA, USA).

## 2 Results and Discussion

### 2.1 Characterization of the tumor-bearing mice

After the tumor cells were inoculated, the growth of subcutaneous tumor masses in the tumor-bearing mice was observed daily.

From day 7, small black and untouchable tumor masses were observed in the B16 melanoma-bearing mice, and the tumor masses grew gradually. The mice were killed on the 15th day, and the tumor masses were weighed. The average tumor mass of the 8 tumor-bearing mice was 0.68 ± 0.15 g. The tumor tissue was stained with hematoxylin and eosin (H&E) for pathological examination, and many tumor cells were observed in the tumor mass (Figure 1).

**Figure 1.**
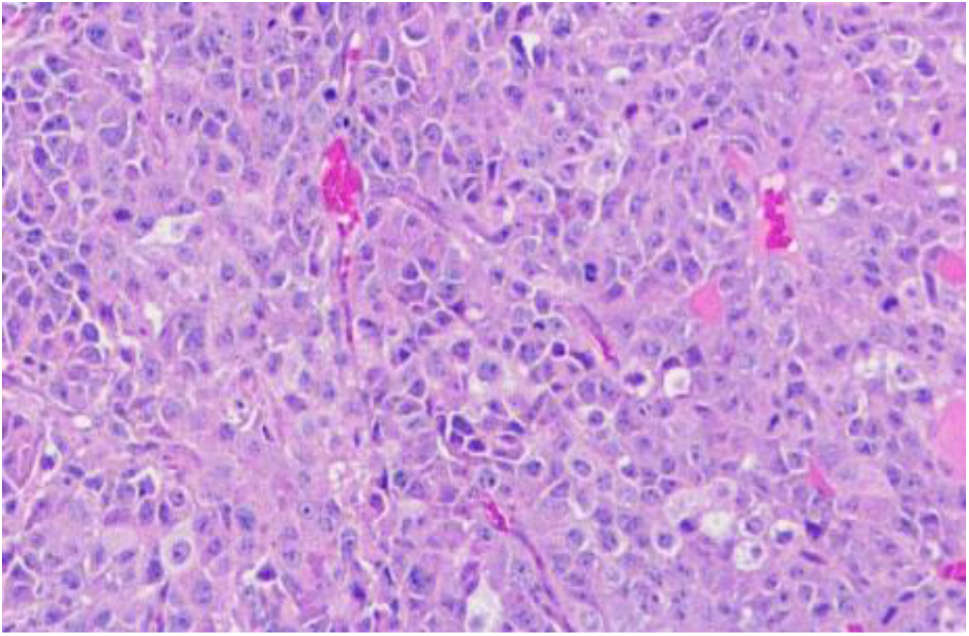
H&E staining of tumors in the B16 tumor-bearing mice (100 ×)

From day 15, small palpable tumor masses were observed in the RM-1 prostate cancer-bearing mice, and the tumor masses grew gradually. The mice were killed on the 23rd day, and the tumor masses were weighed. The average tumor mass of 4 tumor-bearing mice was 0.90 ± 0.13 g. Tumor tissue was stained with hematoxylin and eosin (H&E) for pathological examination, and many tumor cells were observed in the tumor mass (Figure 2).

**Figure 2.**
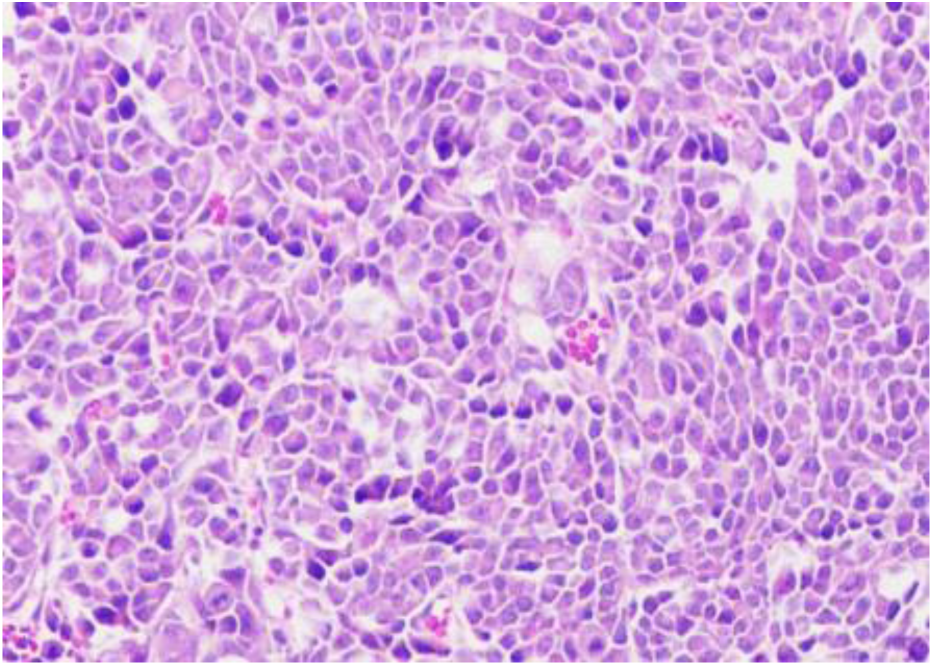
H&E staining of tumors in the RM-1 tumor-bearing mice (100 ×)

### 2.2 Urine proteome changes

Randomly grouped samples at each time point according to different combinations of fold changes (1.5 or 2) and p values (0.01 or 0.05) were used to calculate the average number of differential proteins under different screening conditions. This value was compared with the correct number of differential proteins under this screening condition, and this ratio can be approximated as a false positive rate. The lower the false positive rate, the higher the reliability is. In addition, since the true positive differential proteins are included in the results of all random groupings, the ratio of the approximate false positive rate should be slightly higher than the actual false positive rate (Table 1, Table 2).

**Table 1.**
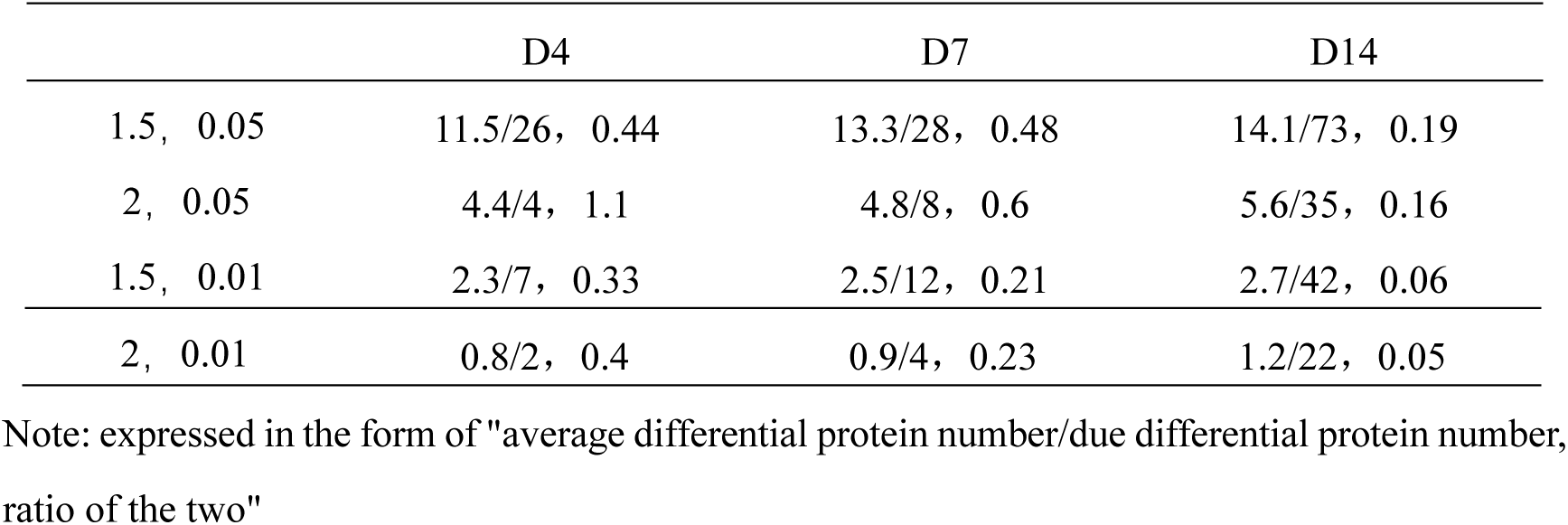
Random grouping results of the B16 tumor-bearing mice under different combinations of p values and fold changes

**Table 2.**
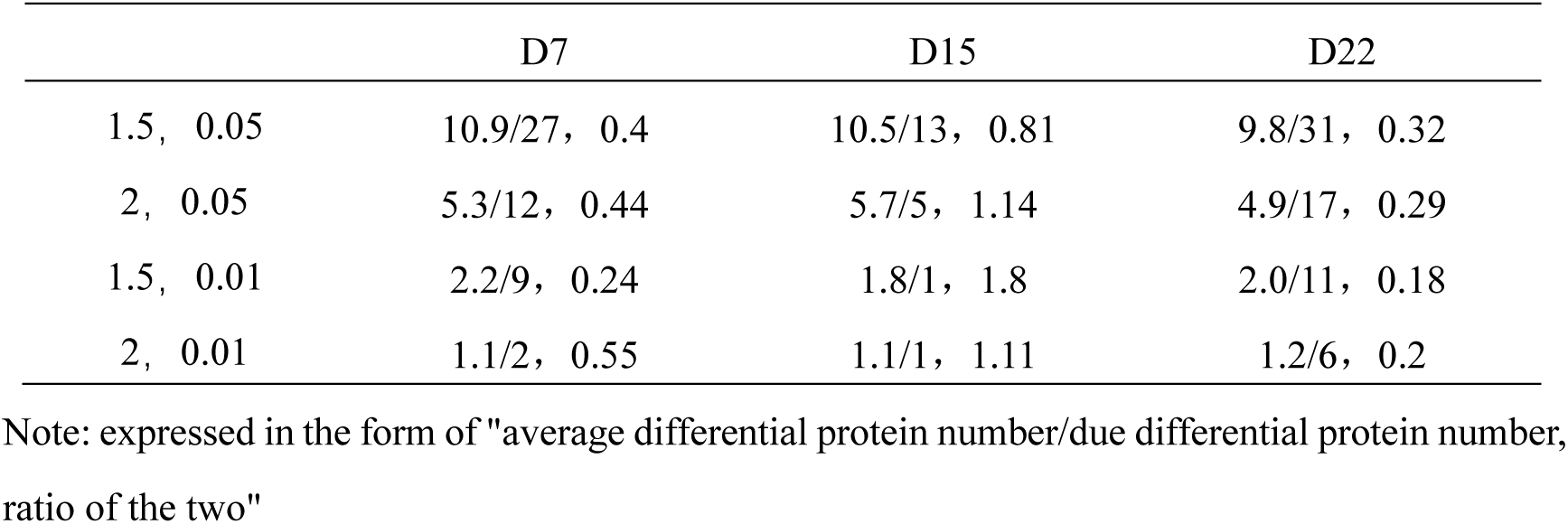
Random grouping results of the RM-1 tumor-bearing mice under different combinations of p values and fold changes

Based on the analysis of the random data of two tumor-bearing models, the screening criteria of the differential proteins we selected were as follows: fold change > 1.5, p value < 0.01. Furthermore, the UniProt database was used to match the human homologous proteins corresponding to these proteins for further analysis.

The differential proteins of the two tumor models at different time points changed significantly with tumor growth, suggesting that the urine proteome has the potential to reflect the occurrence and development of tumors.

#### 2.2.1 B16 melanoma-bearing mice

By DIA quantitative analysis, a total of 532 proteins were identified. Thirty-eight human homologous differential proteins were identified compared with those on Day 0. On Day 4, there were 4 differential proteins, of which 3 were upregulated and 1 was downregulated. On Day 7, there were 7 differential proteins, of which 3 were upregulated and 4 were downregulated. On Day 14, there were 29 differential proteins, of which 21 were upregulated and 8 were downregulated. There were no continuously changing proteins (Figure 3, Table 3).

**Figure 3.**
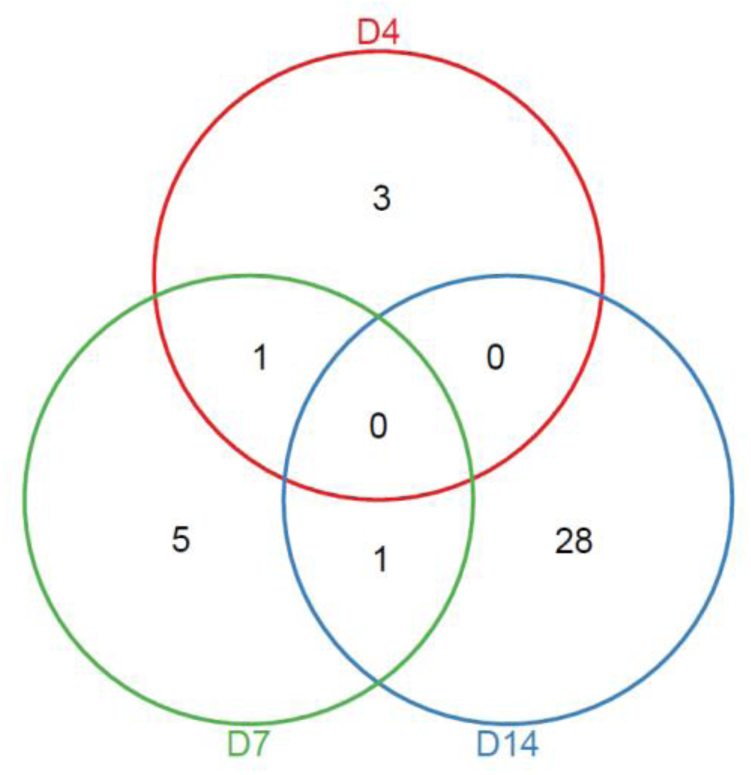
Venn diagram of differential proteins at different stages in the B16 tumor-bearing mice

**Table 3.**
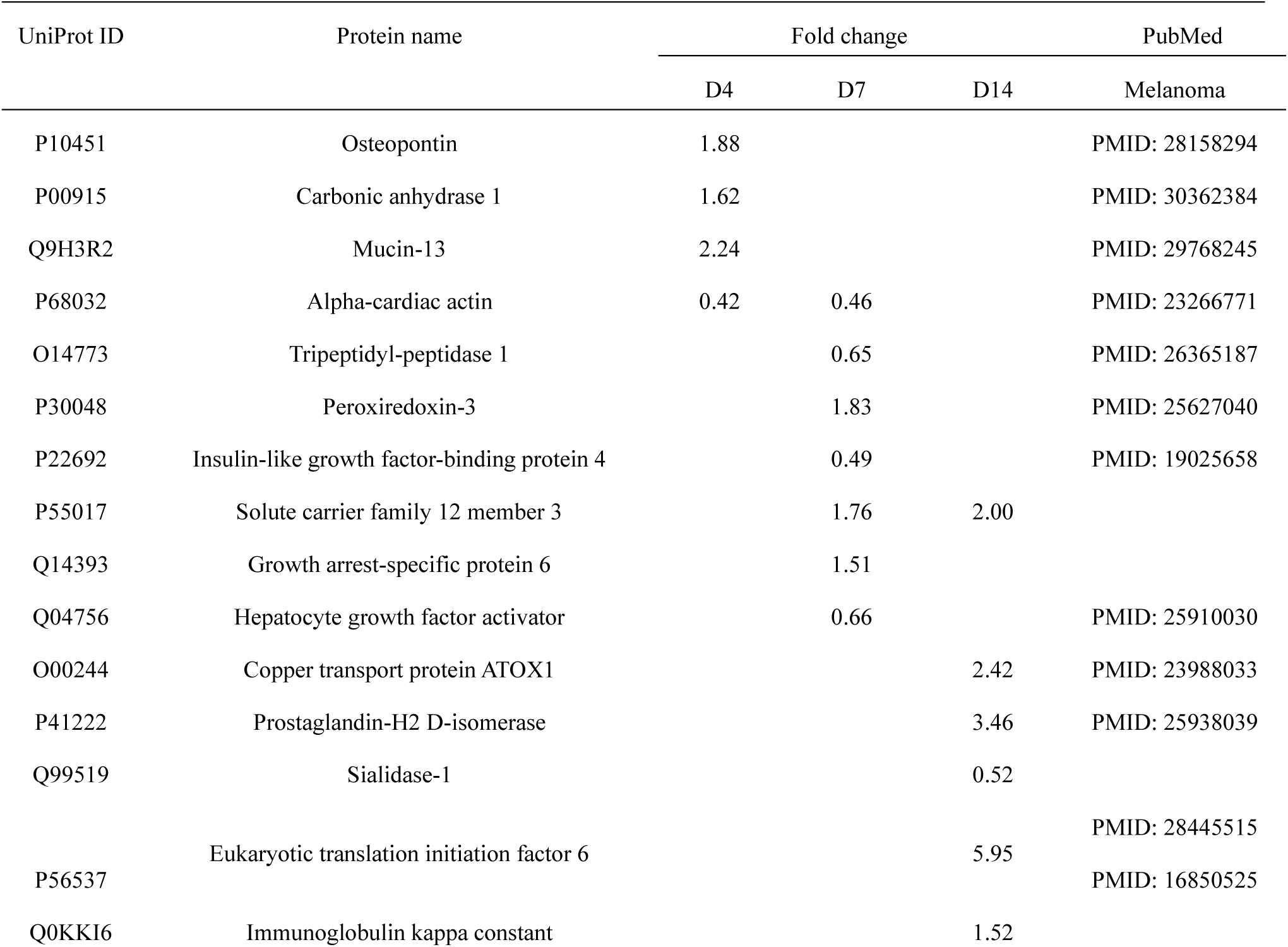
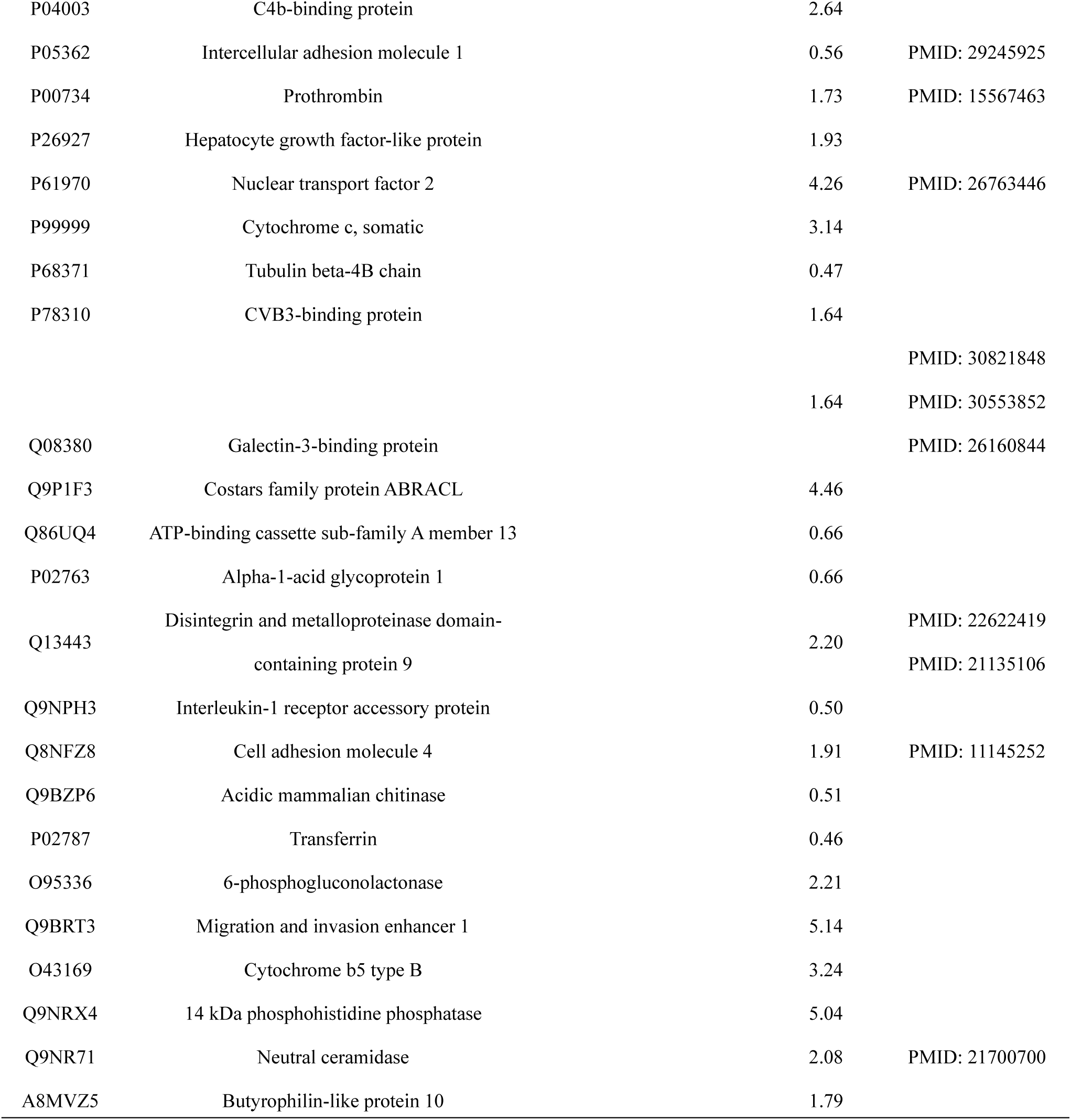
Differential proteins in the B16 tumor-bearing mice

On Day 4, the tumor was not yet visible, and a total of 4 differential proteins were significantly changed, all of which were reported to be related to melanoma. For example, (1) Osteopontin acts as a cytokine and is involved in type I immune responses. Osteopontin-dependent autocrine/paracrine signaling pathways are associated with metastatic melanoma[24], and the protein is involved in cell signaling that regulates tumor progression and metastasis[25]. (2) Carbonic anhydrase 1 participates in interleukin 12-mediated signaling pathways, while Carbonic anhydrase IX inhibitors can increase the sensitivity to chemotherapy and promote the apoptosis of melanoma cells[26]. (3) Mucin-13 is a cell surface glycoprotein that is abnormally expressed in various epithelial cancers. Serum mucin 13 is elevated in 70% of active skin melanoma patients (except for uveal melanoma)[27]. (4) Alpha-cardiac actin is involved in various types of cell movements. Cancer cells need to undergo continuous actin cytoskeleton remodeling to adapt to environmental mechanical stimuli, thereby promoting their survival, migration and metastasis[28, 29]. In addition, β-actin was found to be abnormally expressed in diseases such as melanoma, colorectal cancer, gastric cancer, pancreatic cancer, prostate cancer, and ovarian cancer[30].

On Day 7, the tumor was visible, and a total of 7 differential proteins changed significantly, of which 5 were reported to be associated with melanoma. For example, (1) SPINT2, a proteolytic inhibitor of Hepatocyte growth factor activator, plays an important role in inhibiting the progression of malignant melanoma[31]; (2) downregulation of Insulin-like growth factor-binding protein 4 may be a step in the development of primary melanoma to metastatic melanoma[32]; (3) Peroxiredoxin-3 plays a protective role in cell antioxidant stress. The expression of this protein in the cytoplasm of stromal fibroblasts is related to melanoma-specific survival[33]. (4) Tripeptidyl-peptidase 1 (TPP1) was identified, and POT1-A532P is a POT1 mutation associated with melanoma, which can weaken the interaction of TPP1-POT1[34].

On Day 14, a total of 29 differential proteins were significantly changed, of which 10 were reported to be associated with melanoma. For example, (1) Copper transport protein ATOX1 may be an important cellular antioxidant. The interaction between the anticancer drug cisplatin and the copper transporter ATOX occurs in human melanoma cells[35]. (2) Galectin-3-binding protein has scavenger receptor activity, promotes integrin-mediated cell adhesion, and may stimulate host defense against viruses and tumor cells. Significant upregulation in serum or tumor tissue correlates with poor clinical outcomes in melanoma patients[36–38], and its inhibitors have potential as antimetastatic cancer drugs[39]. (3) Prothrombin plays a role in blood homeostasis, inflammation, and wound healing. Its complex is involved in the production of thrombin in B16F10 melanoma cells[40]; (4) Prostaglandin-H2 D-isomerase may play an important role in the progression of uveal melanoma[41]; (5) Intercellular adhesion molecule 1 is involved in the acute inflammatory response to antigen stimulation. The upregulation of this protein is the key to the lymphatic proliferation of melanoma[42]; (6) Cell adhesion molecule 4 is involved in intercellular adhesion and may have cancer suppressive activity[43]. Melanoma cell adhesion molecule (Mel-CAM) has been shown to be a sensitive marker for epithelioid melanoma [44]; (7) Nuclear transport factor 2 plays an important role in the transport process mediated by the cargo receptor and plays a more general role indirectly. RanBP3 (a potential target for human melanoma therapeutic intervention) regulates melanoma cell proliferation through selective control of nuclear export[45]; (8) Eukaryotic translation initiation factor 6 modulates cell cycle progression and global translation of pre-B cells, and its activation seems to be rate-limiting in tumorigenesis and tumor growth[46] and is positively correlated with tumor tissue content in patients with metastatic melanoma[47]; (9) Disintegrin and metalloproteinase domain-containing protein 9 cleaves and releases many molecules that play important roles in tumorigenesis and angiogenesis and are potential tumor targets[48]. The protein is specifically expressed in stromal fibroblasts and can regulate the proliferation and apoptosis of melanoma cells in vitro and in vivo[49], and its expression is helpful for melanoma invasion[50]; (10) For Neutral ceramidase, acid ceramidase expression regulates the sensitivity of A375 melanoma cells to dacarbazine (the drug of choice for metastatic melanoma)[51].

#### 2.2.2 RM-1 prostate cancer-bearing mice

By DIA quantitative analysis, a total of 406 proteins were identified. Compared with those on Day 0, 14 human homologous differential proteins were identified. On Day 7, there were 9 differential proteins, of which 6 were upregulated and 3 were downregulated. On Day 15, there was no differential protein. On Day 22, there were 6 differential proteins, of which 3 were upregulated and 3 were downregulated. Transferrin changed continuously at the two time points (Figure 4, Table 4).

**Figure 4.**
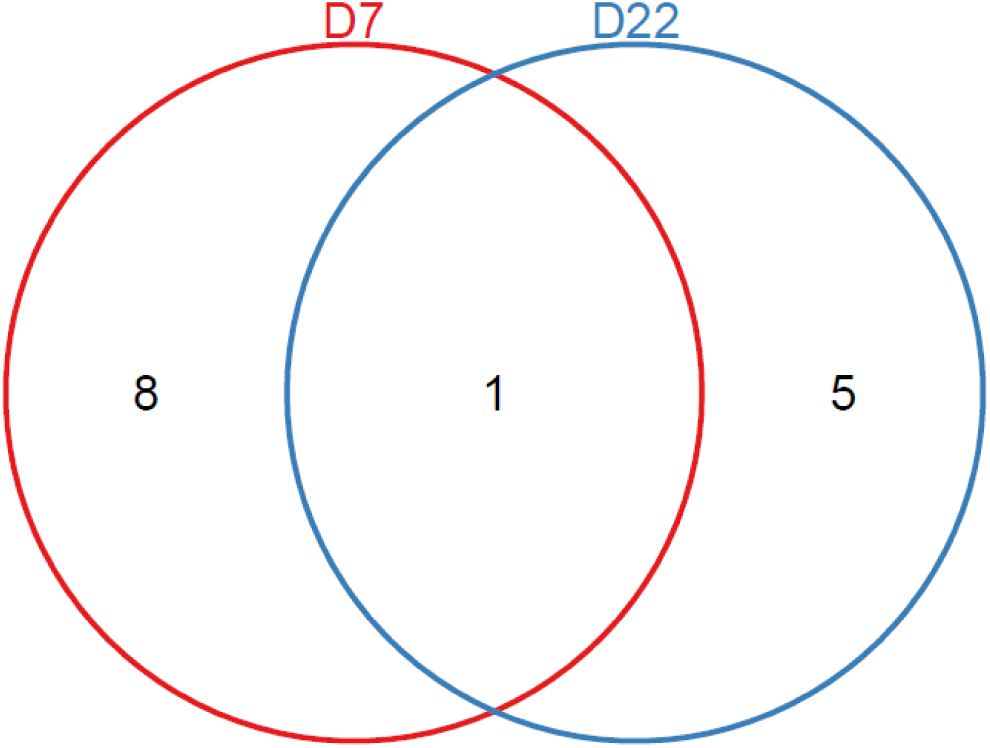
Venn diagram of differential proteins at different stages in the RM-1 tumor-bearing mice

**Table 4.** Differential proteins in the RM-1 tumor-bearing mice

| UniProt ID | Protein name | Fold change |  | PubMed |
| --- | --- | --- | --- | --- |
|  |  | D7 | D22 | Prostate cancer |
| P02749 | Beta-2-glycoprotein 1 | 0.49 |  | PMID: 24367699 |
| P10909 |  |  |  | PMID: 9815554 |
|  | Clusterin | 1.51 |  | PMID: 26313417 |
| Q9Y277 | Voltage-dependent anion-selective channel protein 3 | 1.70 |  | PMID: 25177836 |
|  | Voltage-dependent anion-selective channel |  |  | PMID: 30528266 |
| P21796 | protein 1 | 2.01 |  | PMID: 25177836 |
| Q92896 | Golgi apparatus protein 1 | 1.62 |  |  |
|  |  |  |  | PMID: 2433078 |
|  |  |  |  | PMID: 7029542 |
|  |  |  |  | PMID: 8989919 |
| P02787 |  |  |  | PMID: 28592703 |
|  |  |  |  | PMID: 15619922 |
|  | Transferrin | 0.64 | 0.58 | PMID: 29135121 |
| P04066 | Tissue alpha-L-fucosidase | 1.69 |  |  |
|  |  |  |  | PMID: 29228690 |
| P49747 | Cartilage oligomeric matrix protein | 0.52 |  | PMID: 31413818 |
| O95497 | Pantetheinase | 1.69 |  |  |
| P28799 | Progranulin |  | 1.86 | PMID: 25365768 |
|  | Na(+)/H(+) exchange regulatory cofactor |  |  |  |
| O14745 | NHE-RF1 |  | 0.35 |  |
| P10253 | Lysosomal alpha-glucosidase |  | 1.85 |  |
|  |  |  |  | PMID: 18974881 |
| P54826 | Growth arrest-specific protein 1 |  | 1.81 | PMID: 1505026 |
|  |  |  |  | PMID: 29798970 |
| Q15746 | Myosin light chain kinase, smooth muscle |  | 0.54 | PMID: 19429448 |

On Day 7, the tumor was not yet palpable, and a total of 9 differential proteins changed significantly, of which 6 were reported to be associated with prostate cancer. For example, (1) Clusterin is upregulated in prostate cancer solid tumors [52], and the intracellular level in patients is related to the Gleason score[53]; (2) Transferrin is an iron-binding transporter and an effective mitogen for prostate cancer[54, 55], and upregulation of Transferrin is often associated with increased adhesion, invasion, and metastasis[56]. For example, Transferrin receptors are direct MYC target genes, and MYC oncogenes are important mediators of tumor initiation and progression in prostate cancer[57]. Studies have shown that the Transferrin concentration in the prostate fluid of prostate cancer patients is significantly higher than that of patients with benign prostatic hyperplasia and normal controls[58]. Compared with the healthy controls, 18/22 prostate cancer patients had a significant increase in Transferrin in their urine[59]. (3) Voltage-dependent anion-selective channel protein 1 participates in the formation of channels across the mitochondrial outer membrane and plasma membrane. The mRNA expression of its genes is significantly increased in prostate cancer tissues [60]. Zinc and p53 destroy the mitochondrial binding of hexokinase 2, which has dual metabolic and apoptotic functions in prostate cancer cells through phosphorylation of voltage-dependent anion selective channel protein 1[61]; (4) Voltage-dependent anion-selective channel protein 3 participates in the formation of a channel across the outer mitochondrial membrane, and there is no significant difference in the mRNA expression of its genes in prostate cancer tissues[60]; (5) Cartilage oligomeric matrix protein promotes the progression of prostate cancer by enhancing infiltration and disrupting intracellular calcium homeostasis[62]; this protein in patients with osteoarthritis is independently associated with metastatic disease of prostate cancer [63]. Studies have shown that prostate cancer cells tend to show bone metastasis, which is the main cause of morbidity and mortality in prostate cancer patients[64]. (6) Beta-2-glycoprotein 1 may prevent activation of the intrinsic coagulation cascade by binding to phospholipids on the surface of damaged cells. Human phosphatidylserine-targeting antibody fragments can be used as highly specific tumor imaging agents by interacting with Beta-2-glycoprotein 1 to form high-affinity complexes. This study has been verified in nude mice with subcutaneous or in situ human prostate tumors[65].

On Day 15, the tumor was just palpable, and there was no significant change in protein.

On Day 22, a total of 6 differential proteins were significantly changed, of which 4 were reported to be associated with prostate cancer. For example, (1) Progranulin is a growth factor involved in inflammation[66], tumorigenesis and development[67] and may play a key role in the progression of prostate cancer, promoting movement and proliferation [68]; (2) Growth arrest-specific protein 1 is a growth inhibitory protein that participates in growth inhibition and prevents entry into S phase [69], and 8 gene tags including this gene have the potential to diagnose prostate cancer[70]; (3) Myosin light chain kinase in smooth muscle participates in inflammatory response, promotes cell migration and tumor metastasis and is a new downstream target of the androgen signaling pathway in prostate cells. Androgen can downregulate the expression of myosin light chain kinase in LNCaP human prostate cancer cell lines[71], and circRNA-myosin light chain kinase can promote the progression of prostate cancer[72].

At present, it is not clear why this phenomenon occurred on Day 15 without differential proteins. By reviewing the original data, we found that the condition that limited the screening of differential proteins was mainly the p value. At this time point, there were only five human homologous differential proteins under the screening condition of “fold change greater than 1.5 times and p value less than 0.05”: Acid sphingomyelinase-like phosphodiesterase 3b, Acid sphingomyelinase-like phosphodiesterase 3a, Galectin-3-binding protein, Adhesion G-protein coupled receptor G1, and Cartilage oligomeric matrix protein. These five proteins are mainly involved in lipid metabolism, cell adhesion, and the immune response and participate in various signal transduction pathways, which are involved in tumorigenesis; that is, when the body fights tumor cells, tumor cells prepare for subsequent infinite growth, invasion and metastasis.

#### 2.2.3 Comparison of the differential proteins in the urine of the B16 melanoma, RM-1 prostate cancer and Walker 256 tumor-bearing models

In total, 37 and 13 unique differential proteins were identified (Figure 5) by comparing B16 melanoma and RM-1 prostate cancer in the tumor-bearing mice, and the only overlapping protein was Transferrin. It has been reported that Transferrin is widely used as a tumor-targeting ligand for anticancer drugs because the transferrin receptor is overexpressed on the surface of various rapidly growing cancer cells, which have an affinity for Transferrin that is 10-100 times higher than that of normal cells, thus providing an effective means of targeting cancer cells for treatment[73, 74]. Of the above unique differential proteins, 17/37 have been reported to be related to melanoma, and 8/13 have been reported to be related to prostate cancer; that is, the urine proteome can reflect the differences in the production of different tumors in the same animals and at the same site. Presumably, the reason for the substantial difference in the number of proteins between the two models is that the subcutaneous tumor-bearing model of melanoma is closer to the in situ model, which has a greater impact on the urine proteome.

**Figure 5.**
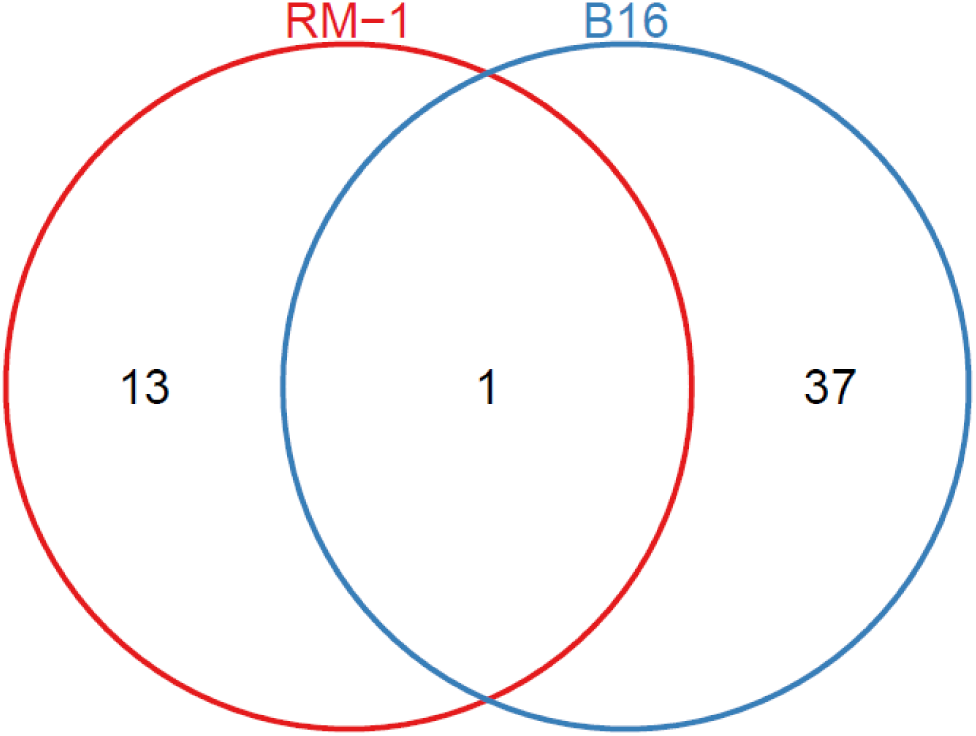
Venn diagram of differential proteins between the B16 and RM-1 tumor-bearing models

At p < 0.01, comparing homologous proteins among the two tumor models of B16 and RM-1 model and laboratory research on a Walker 256 (W256) carcinosarcoma model[16], we found distinctive differential proteins (31, 12, and 49), all of which comprised more than half of each differential protein type. There was no common variation in the differential proteins in the three tumor-bearing animal models, but there were common differences between every two models (Figure 6). Overlapping proteins are 6-phosphate glucose acid esterase, intercellular adhesion molecule 1, nuclear transport factor 2, carbonic anhydrase 1, galactosyl lectin-3-binding protein, lysosome alpha glycosidase enzymes, transferrin, and acidic mammalian chitinase. The molecular functions and biological processes involved in the above 8 proteins are mostly related to energy metabolism, material transport, inflammatory response and immune response. To some extent, the results of comparisons showed that there was a certain similarity in the growth of different tumors at the same site, but the difference reflected in the urine proteomes was much greater than the similarity; that is, the urine proteome was sensitive enough to reflect the changes of different tumors.

**Figure 6.**
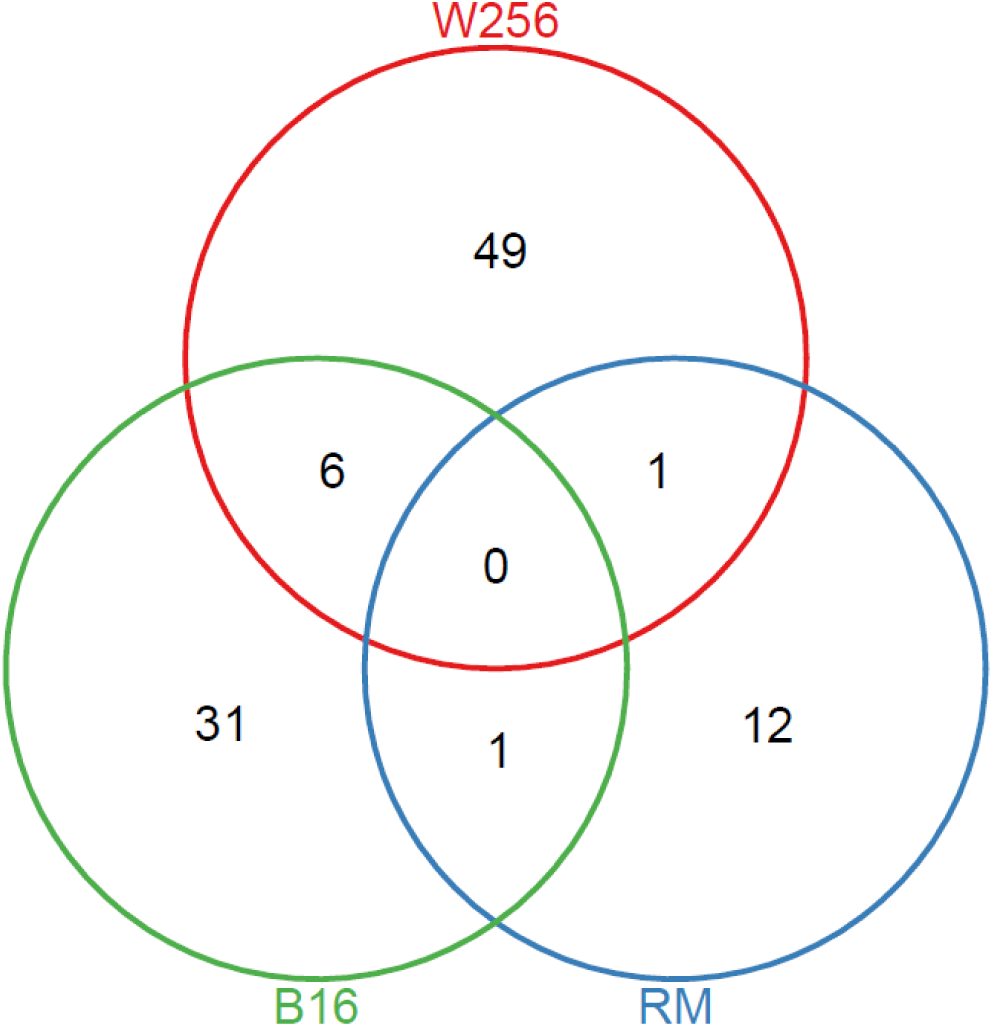
Venn diagram of differential proteins among the B16, RM-1 and W256 tumor-bearing models

A comparison was performed with the differential proteins obtained from three tumor-bearing animal models (B16, RM-1 and W256) in the early stage (before the tumor was detectable) (Figure 7). We found that the three tumor-bearing animal models had no differential proteins with common changes, and there were no differential proteins between every two models; that is, urine proteomics is sensitive enough to reflect changes in the growth of different tumors, even before the tumor is detectable.

**Figure 7.**
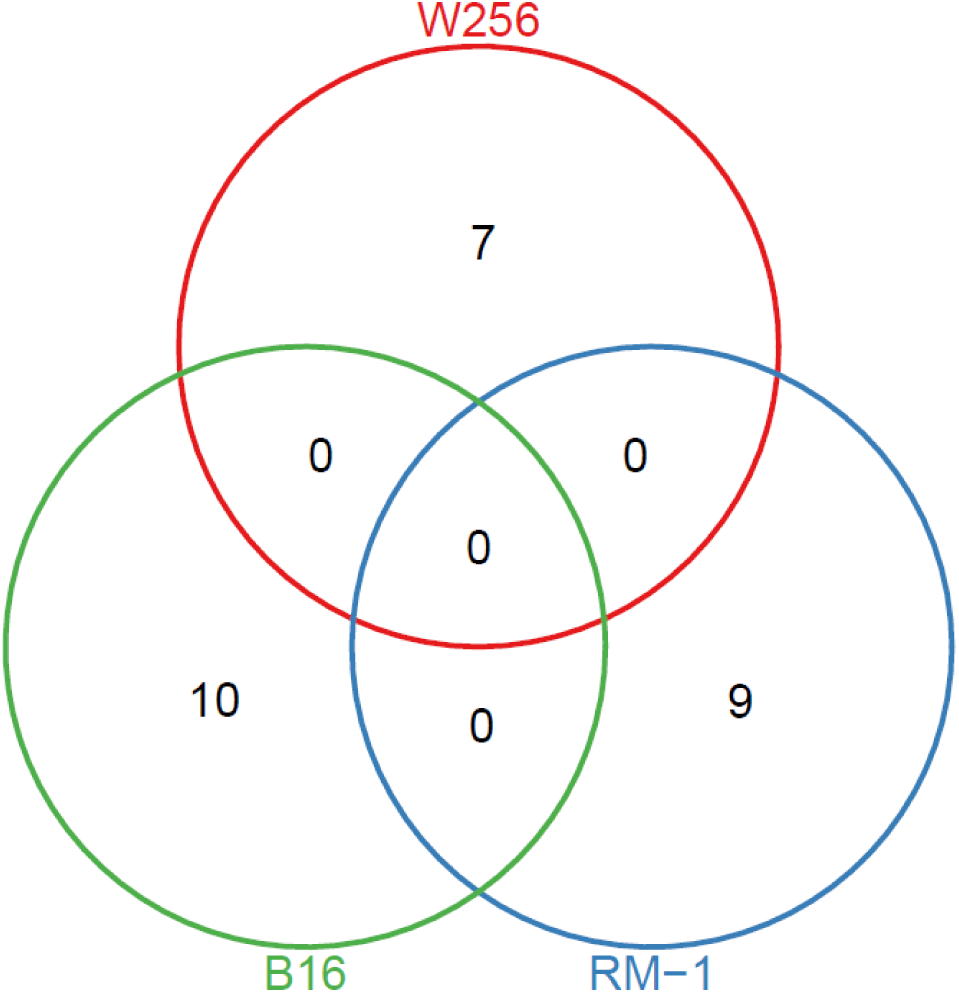
Venn diagram of early differential proteins among the B16, RM-1 and W256 tumor-bearing models

The above results suggest that urine has potential for early diagnosis, differentiation of different tumor types and development.

### 2.3 Functional enrichment analysis of the differential proteins

DAVID was used to annotate the function of the differential proteins, showing the biological process, cell composition and molecular function of these proteins at each time point. Pathway analysis of the proteins was performed by using IPA.

In general, the more stringent the screening conditions are, the lower the average differential protein number, the number of expected differential proteins, and the false positive rate are. However, the more stringent the screening conditions are, the fewer differential proteins will be obtained, and the false-negative proteins will be screened out. When the differential proteins obtained under strict screening conditions are used for biological function analysis, there will be less relevant information. Therefore, in the biological function analysis of the differential proteins, this experiment mainly analyzed the proteins screened with the condition of “fold change > 1.5, p value < 0.01”, which supplemented the proteins screened under the condition of “fold change > 1.5, p value < 0.05”, for reference.

#### 2.3.1 B16 melanoma-bearing mice

According to the analysis of biological processes, apoptotic processes were shown on Day 7, and cell adhesion and inflammatory processes were shown on Day 14, which were related to tumor invasion and metastasis (Figure 8A). Analysis of cell composition revealed that the differential proteins were secreted proteins, mostly from the extracellular matrix, cell surface and membrane (Figure 8B). The molecular functions were similarly analyzed, and from Day 7 onwards, the functions involved in apoptosis, binding and transport are shown (Figure 8C). The analysis of pathways revealed that the pathways at different time points had their own characteristics, and some of them were reported to be related to melanoma (Table 5). For example, before the tumor can be observed, the RhoA signal was significantly enriched. RhoA is reported to be the main signal transduction factor activated by GPR56 in melanoma cells, thus promoting cell movement[75]. CMSP induces the differentiation of B16F1 melanoma cells through the RhoA-MARK signaling pathway[76]. The crosstalk between dendritic cells and natural killer cells may represent the mechanism by which growing tumors evade immune responses. This interaction has an important effect on the immune response of pathogens and potential tumor cells, as demonstrated in melanoma cells[77]. The FAK signal can inhibit the migration and metastasis of B16F10 cells, which may be a potential target for the treatment of melanoma[78]. Focal adhesion kinase (FAK) is a nonreceptor tyrosine kinase that is involved in tumor angiogenesis. In addition to its role as a kinase, FAK is involved in cell proliferation, adhesion, migration, invasion and apoptosis[79–82]; however, acute response signals, LXR/RXR activation, and FXR/RXR activation were significantly enriched after the tumor was observed. It has been reported that FXR/RXR activation may attempt to reduce inflammation, usually in conjunction with LXR and other receptors, to activate homeostasis in cholesterol and triglyceride metabolism and inhibit inflammation[83].

**Figure 8A.**
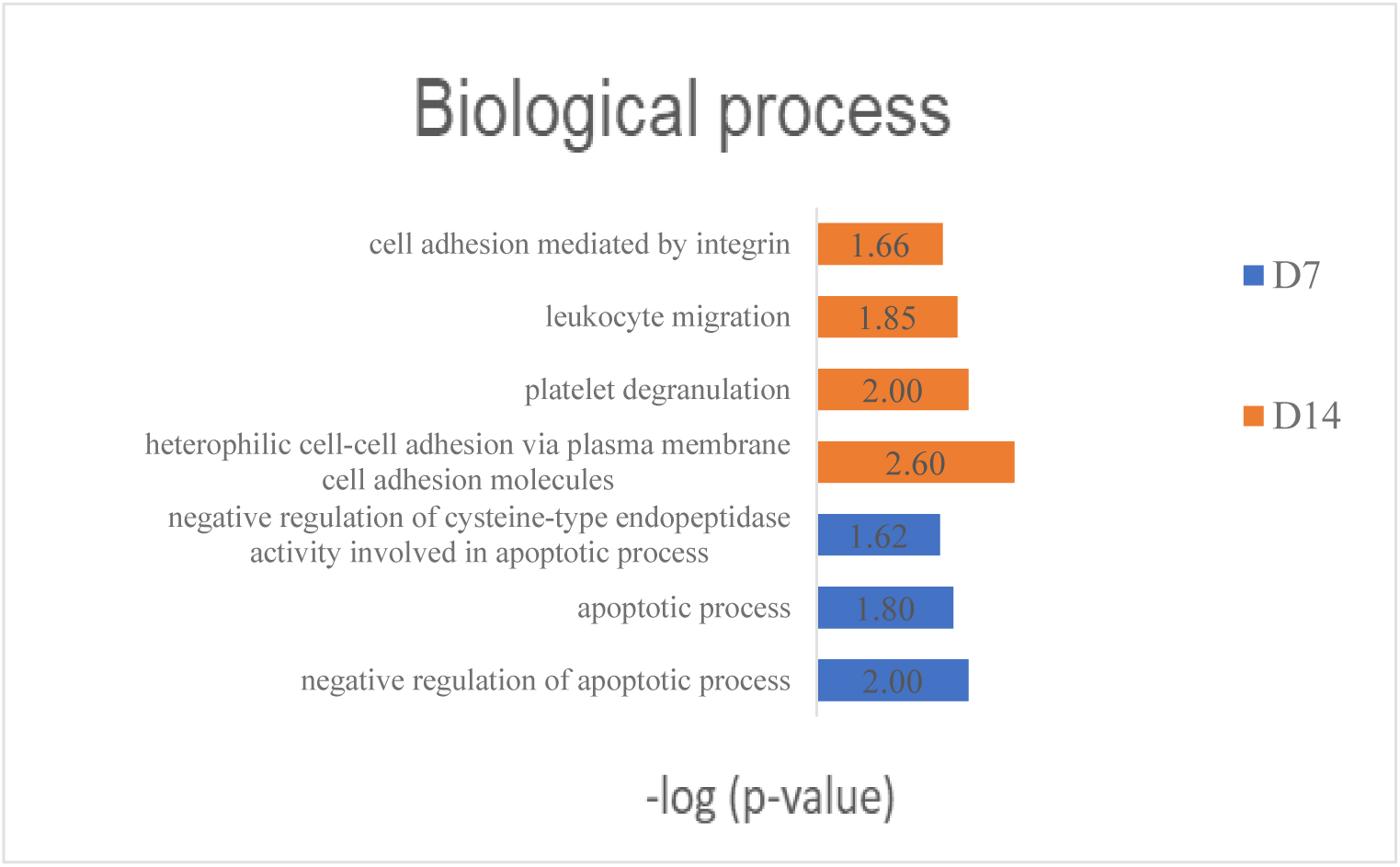
Biological process analysis of differential proteins in the B16 tumor-bearing mice

**Figure 8B.**
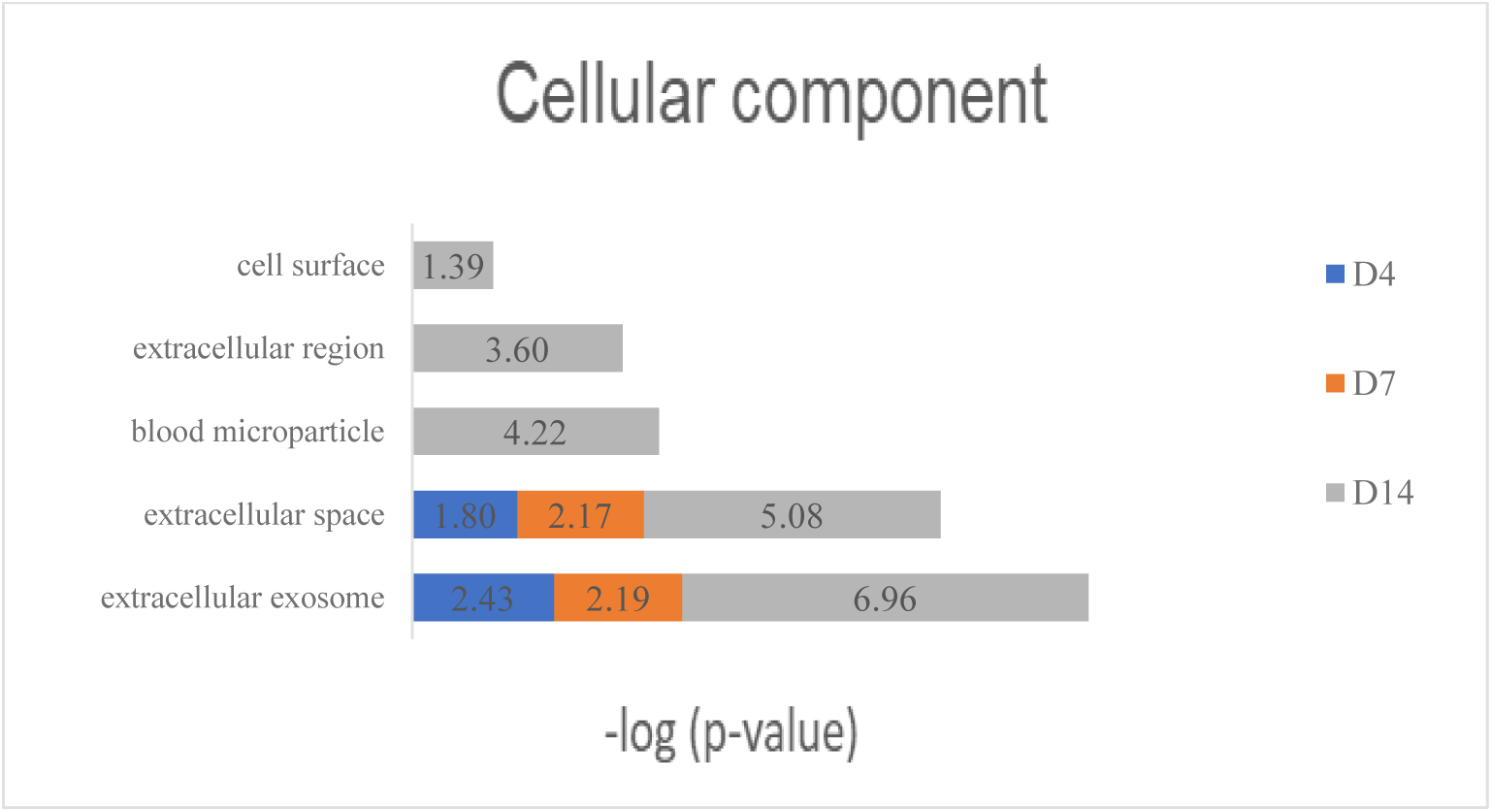
Cell component analysis of differential proteins in the B16 tumor-bearing mice

**Figure 8C.**
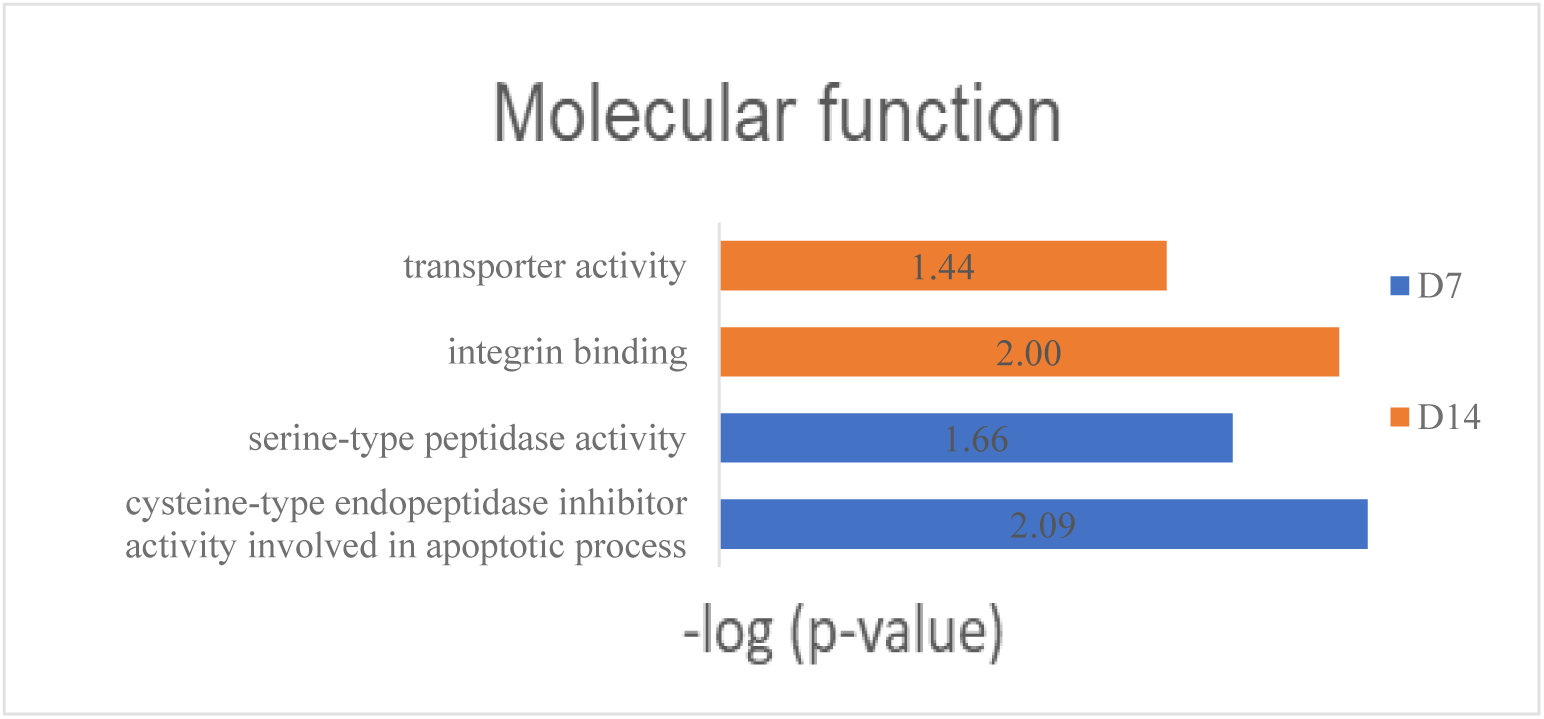
Molecular function analysis of differential proteins in the B16 tumor-bearing mice

**Table 5.**
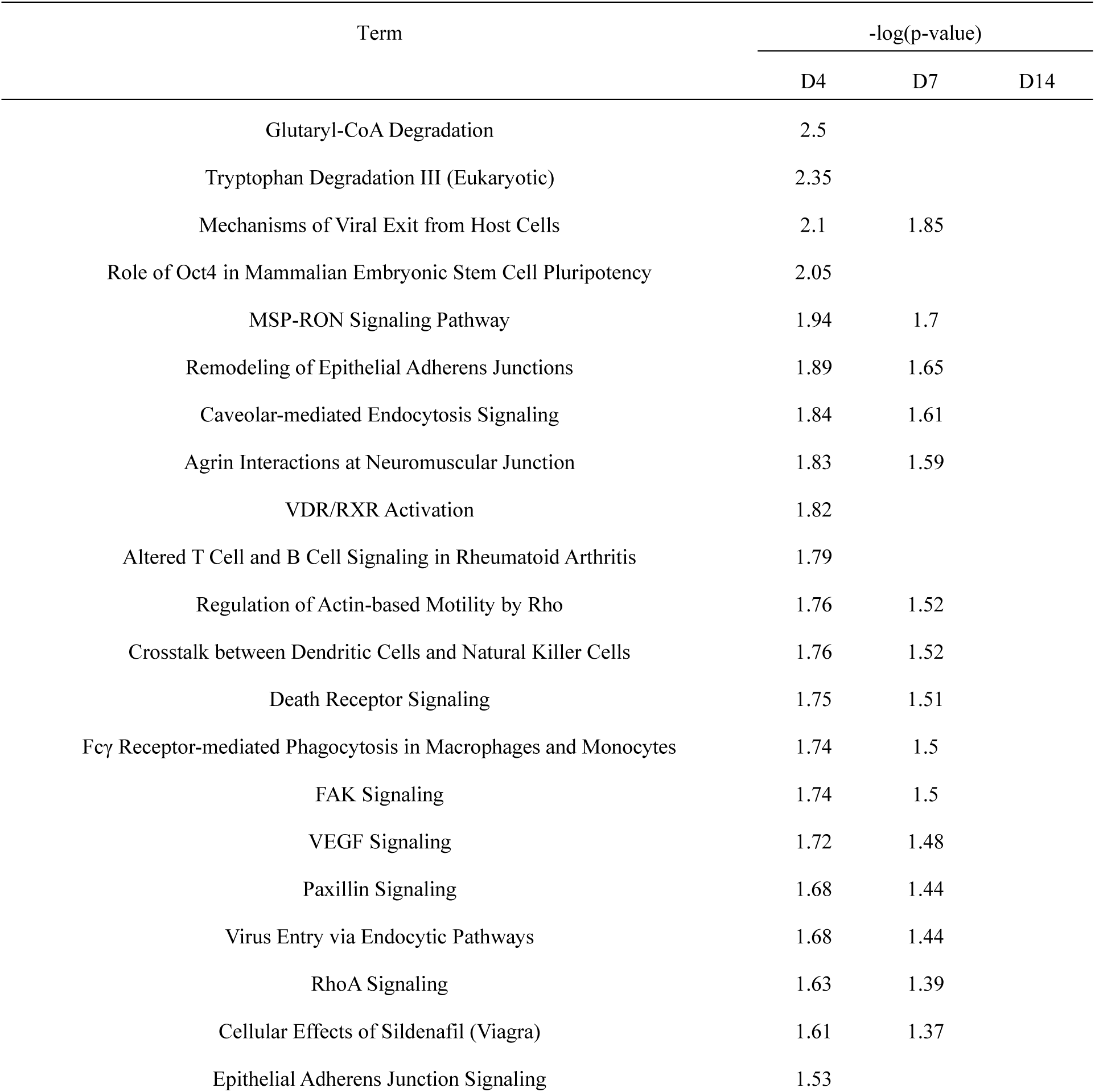
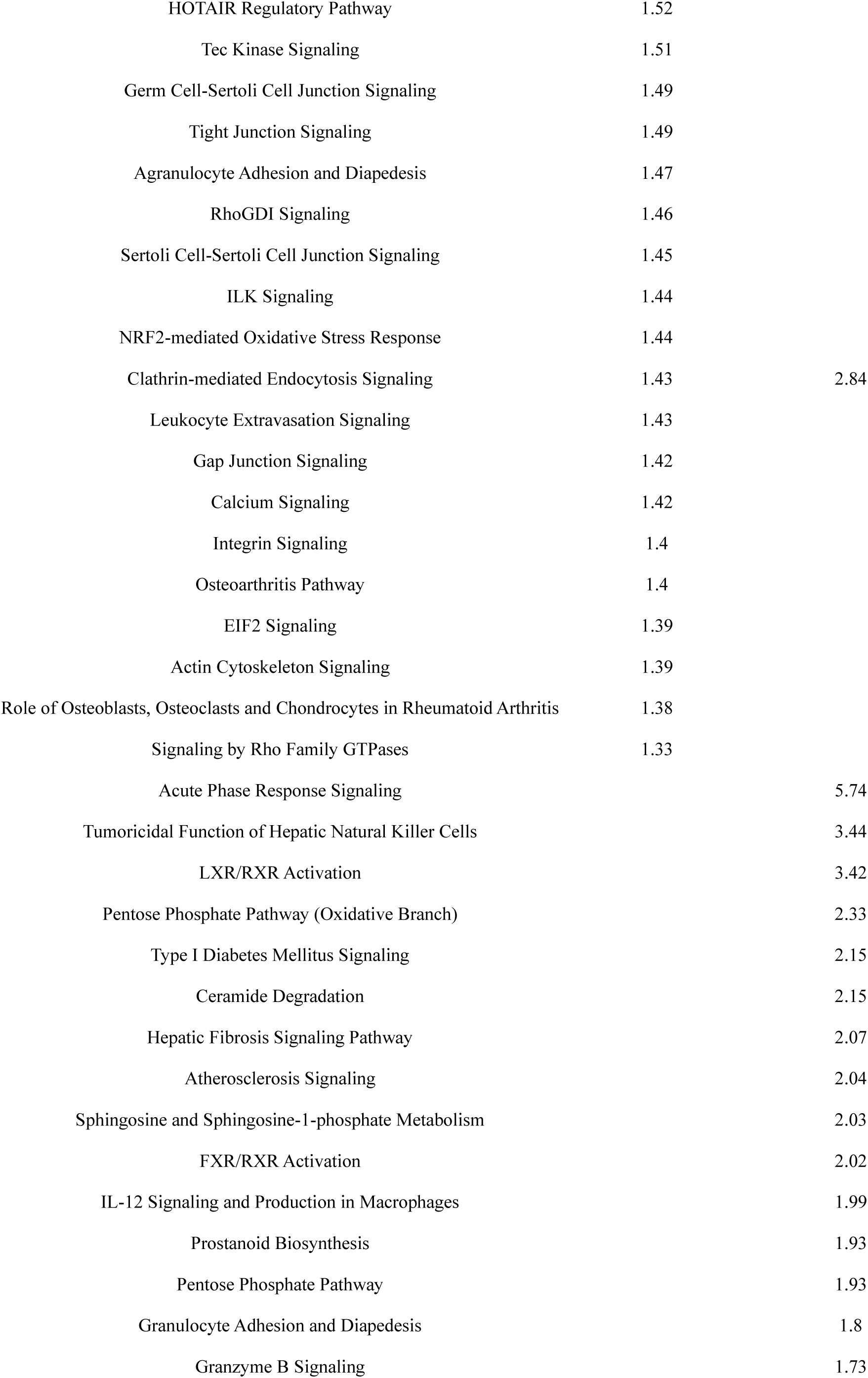
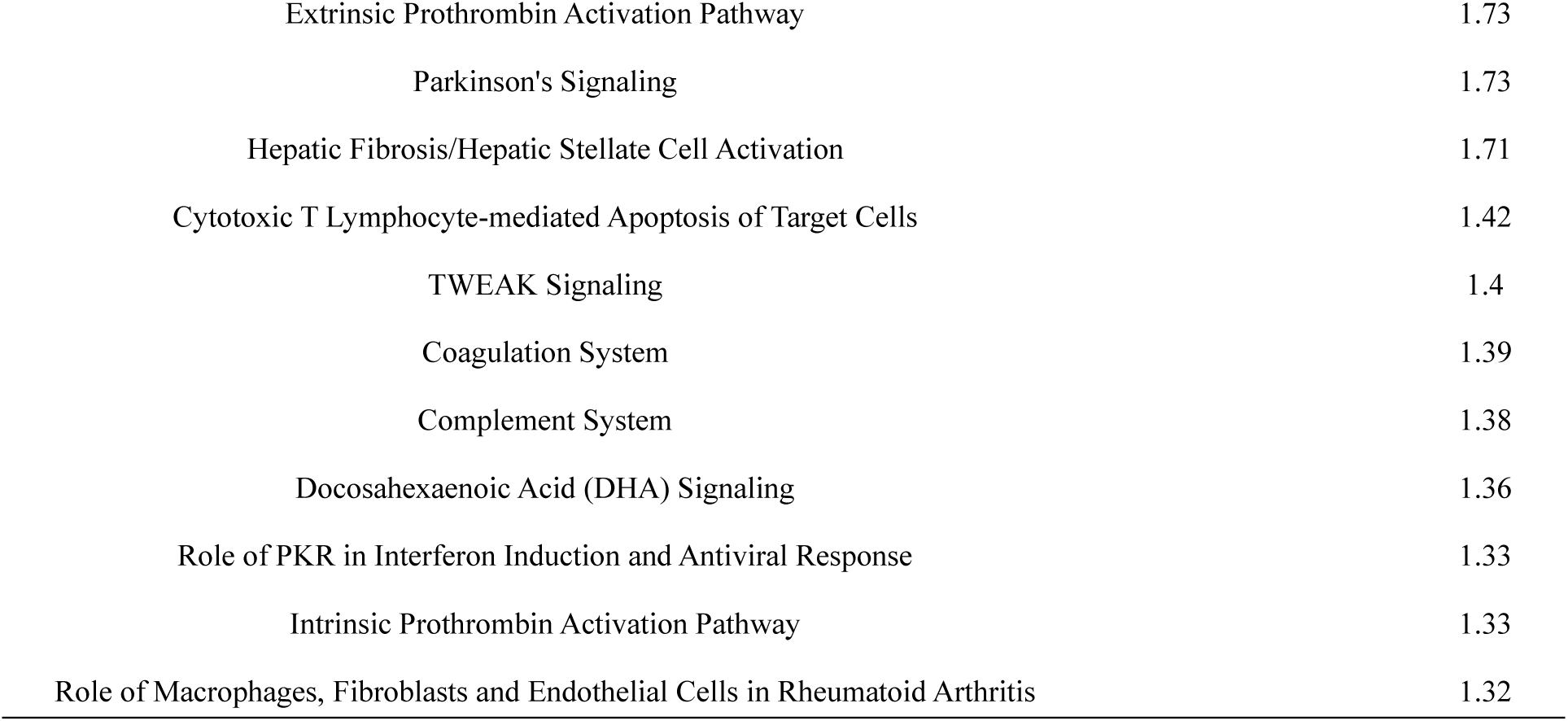
Pathway analysis of differential proteins in the B16 tumor-bearing mice

A total of 86 homologous differential proteins were identified with the condition of “fold change > 1.5, p value < 0.05”, among which 39 differential proteins were reported to be associated with melanoma (Table 6, Figure 9).

**Figure 9A.**
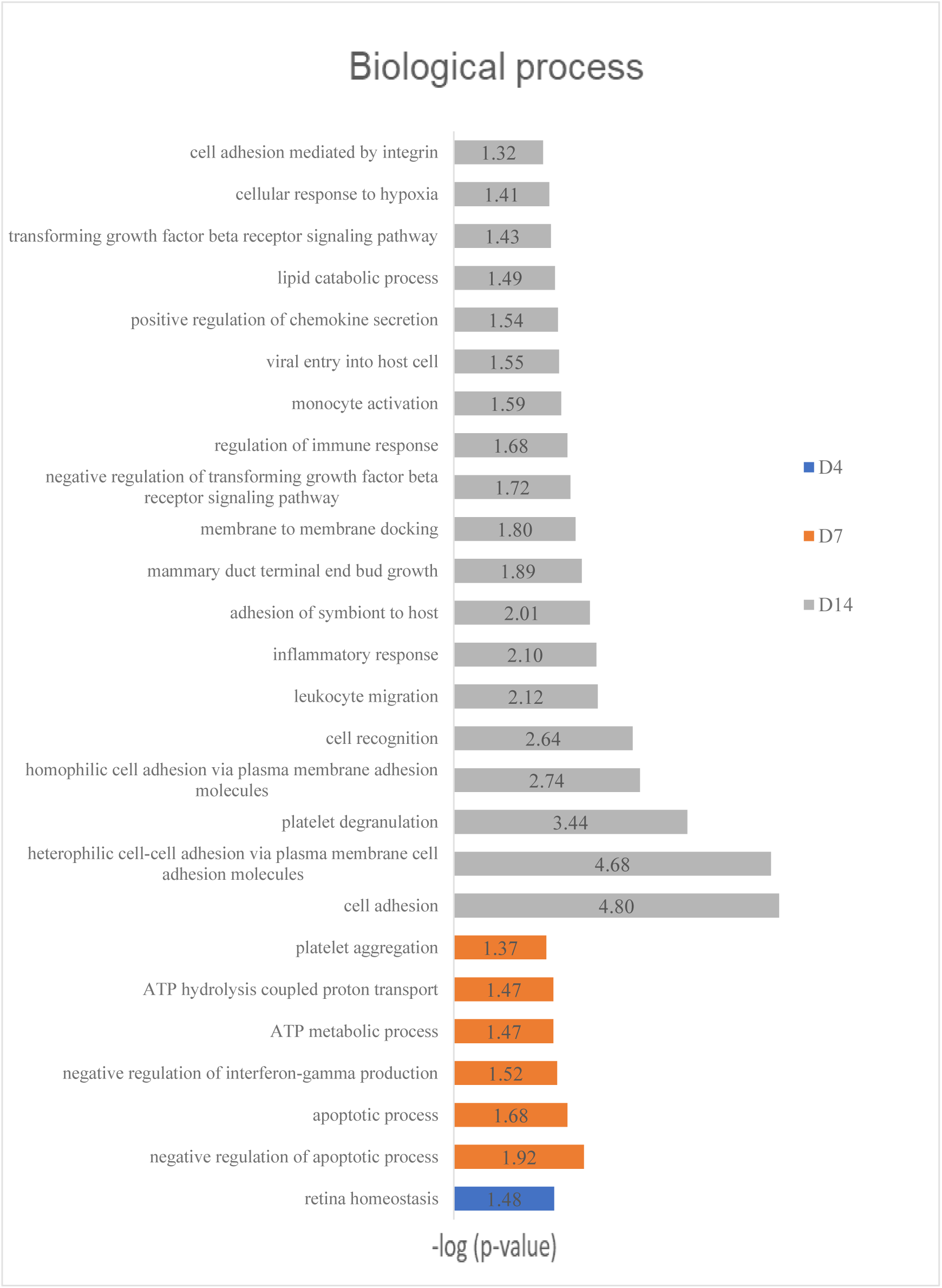
Biological process analysis of differential proteins in the B16 tumor-bearing mice (p<0.05)

**Figure 9B.**
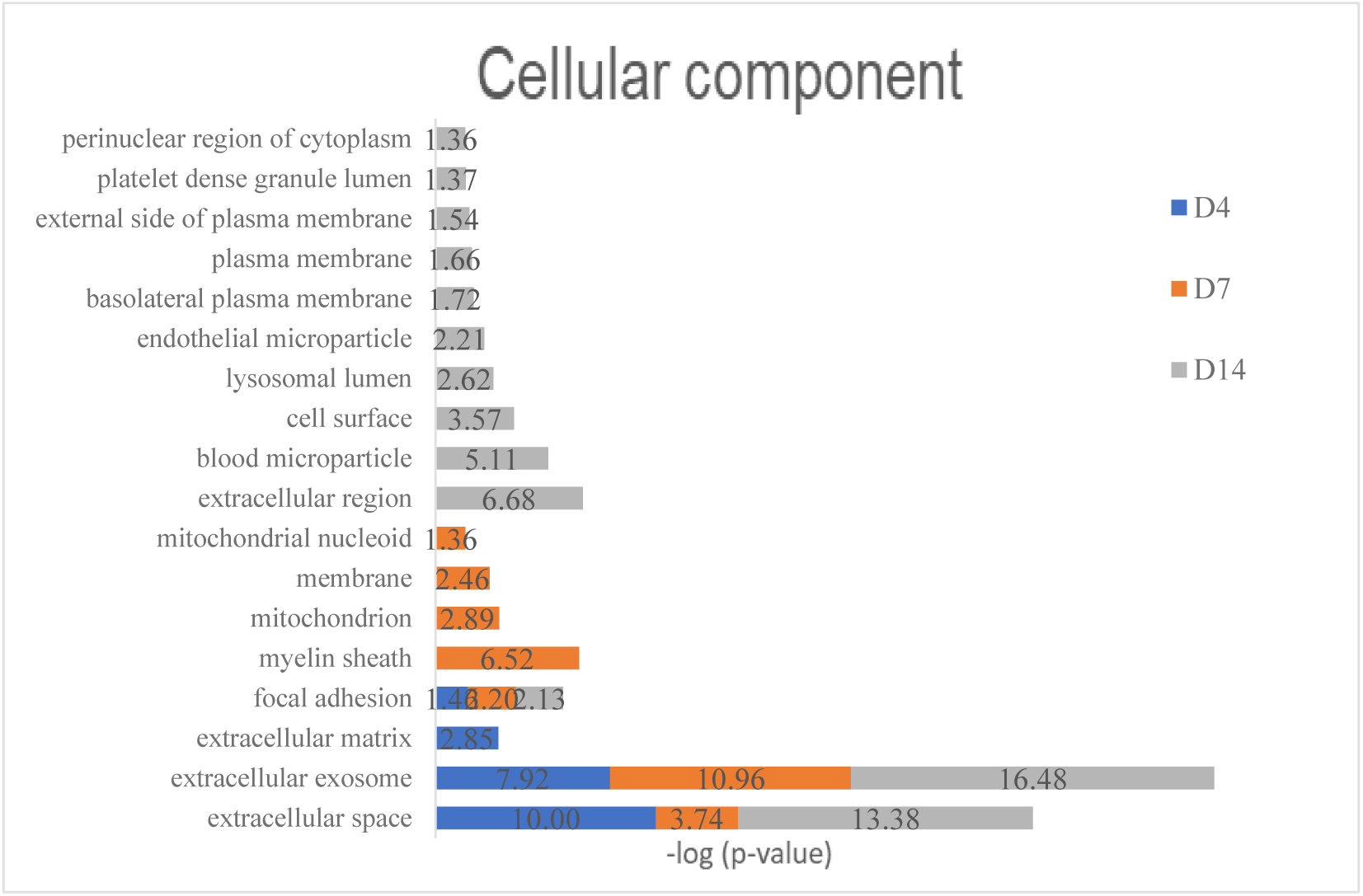
Cell component analysis of differential proteins in the B16 tumor-bearing mice (p<0.05)

**Figure 9C.**
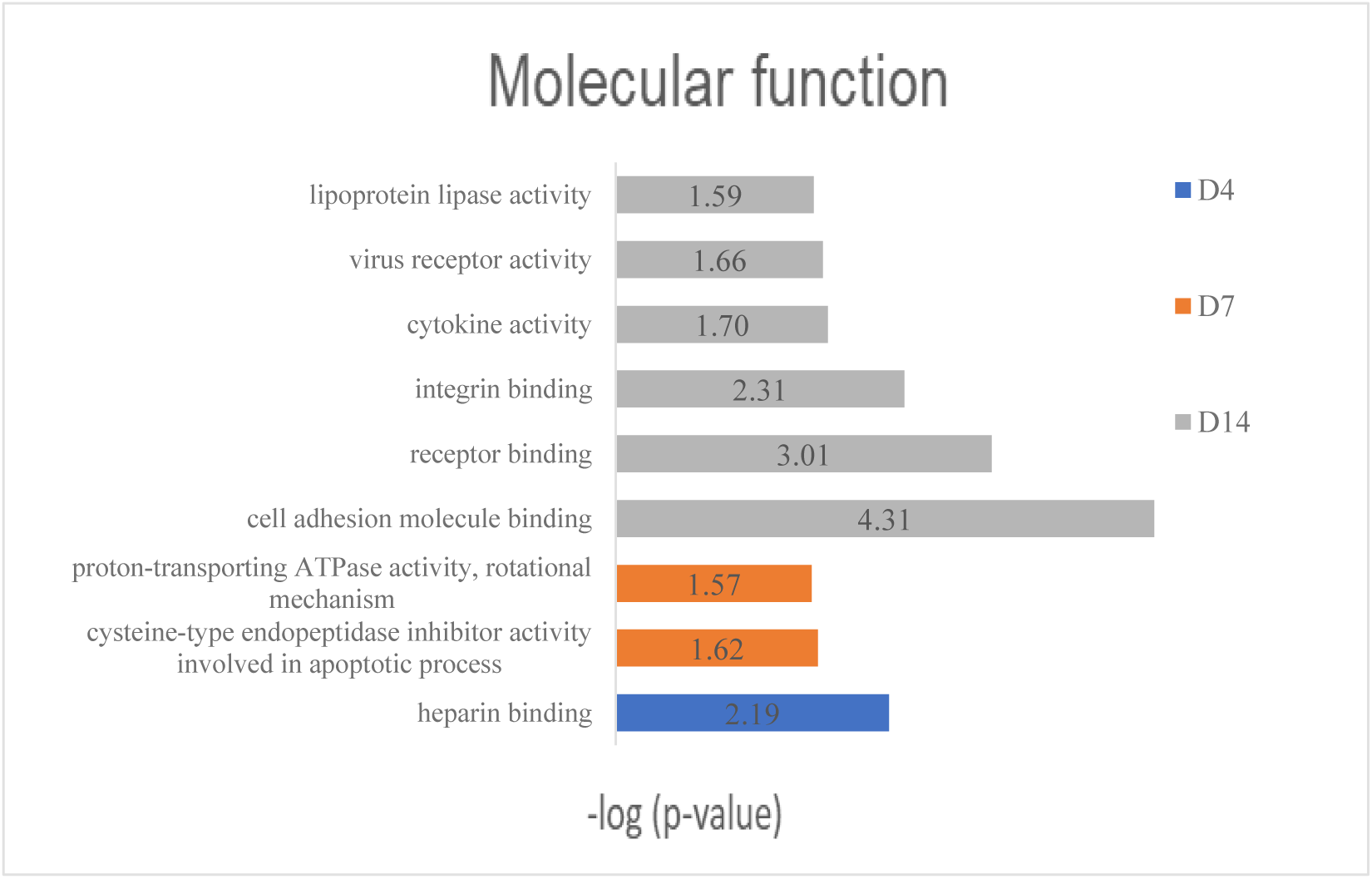
Molecular function analysis of differential proteins in the B16 tumor-bearing mice (p<0.05)

**Table 6A.**
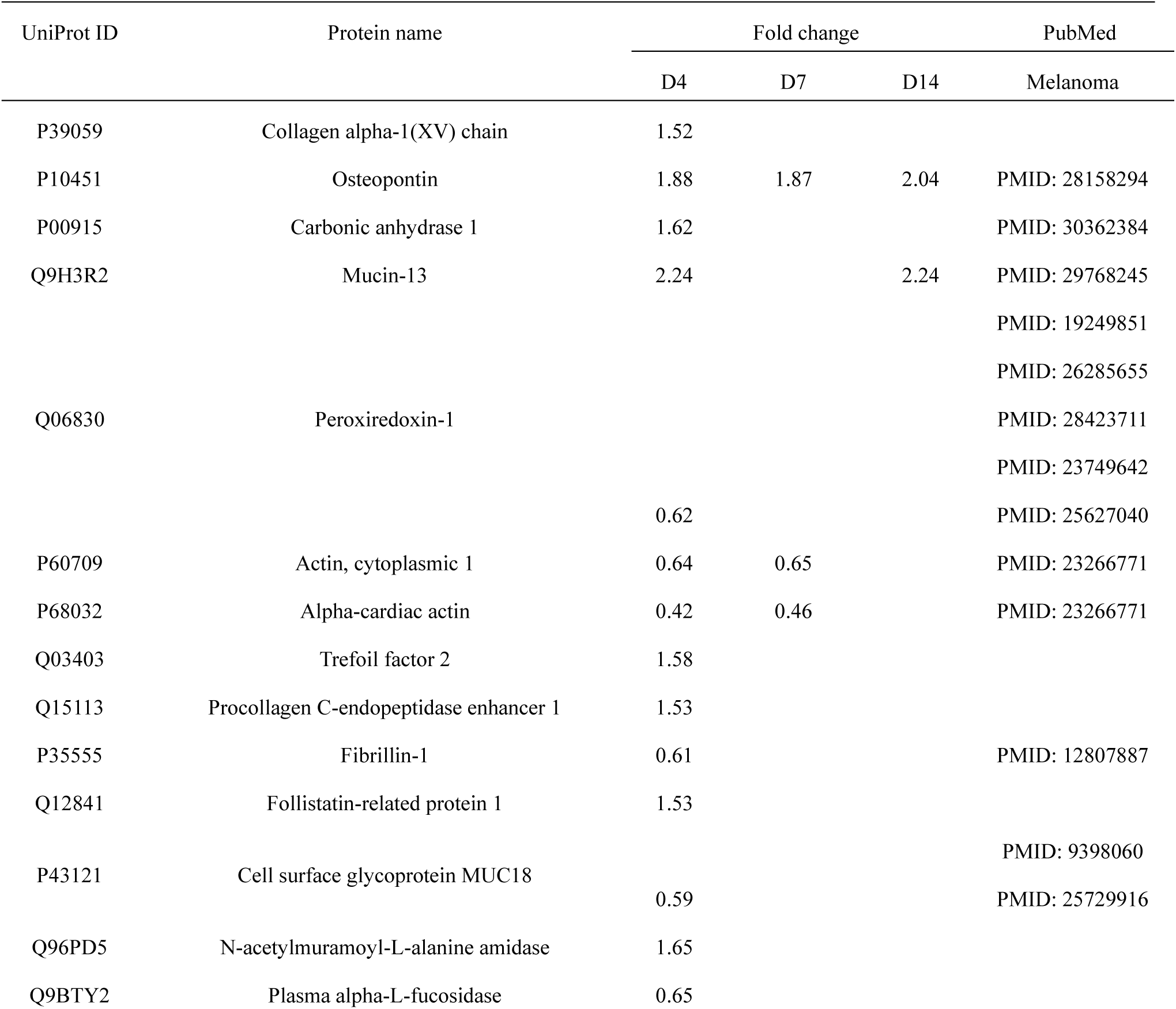
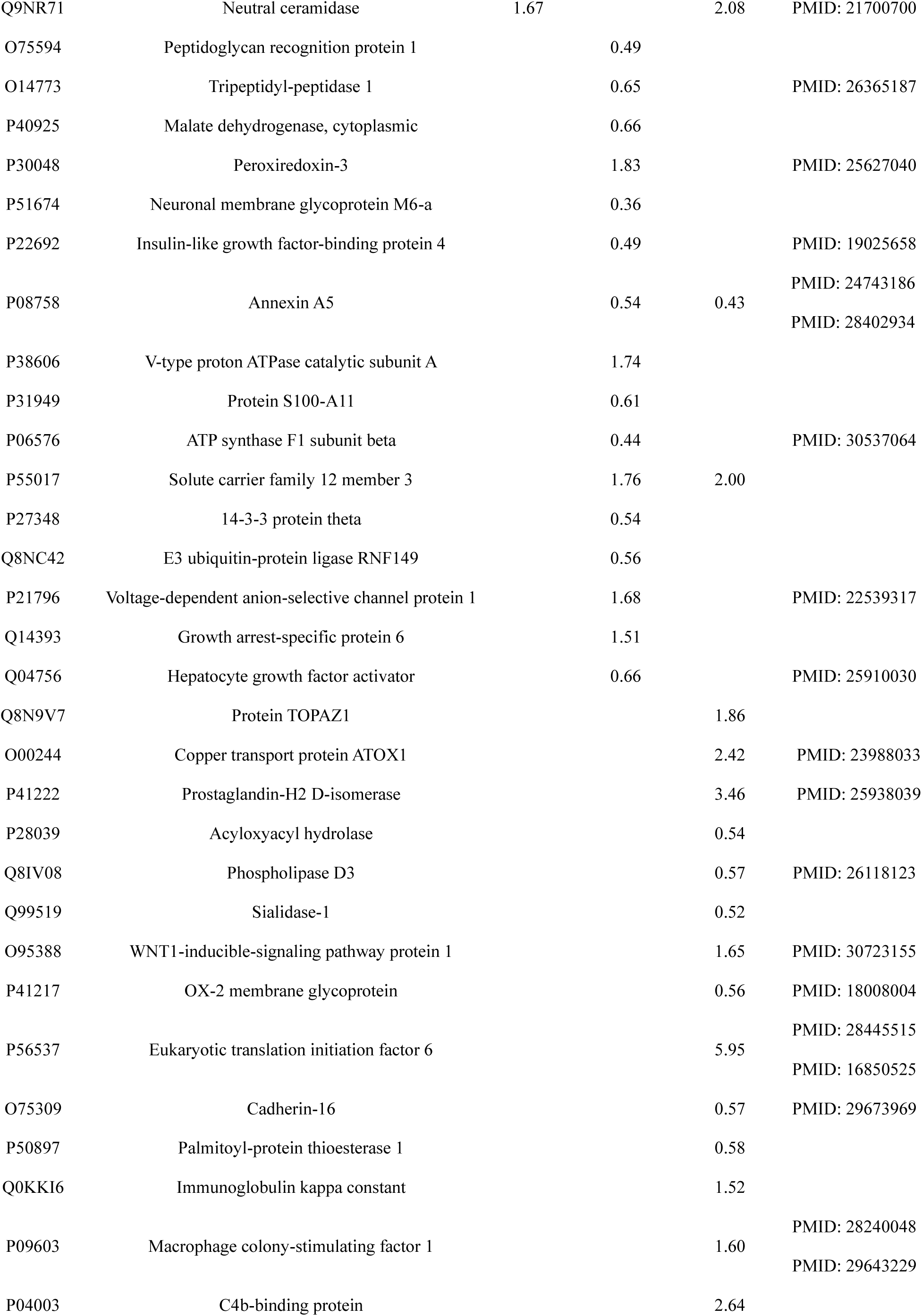
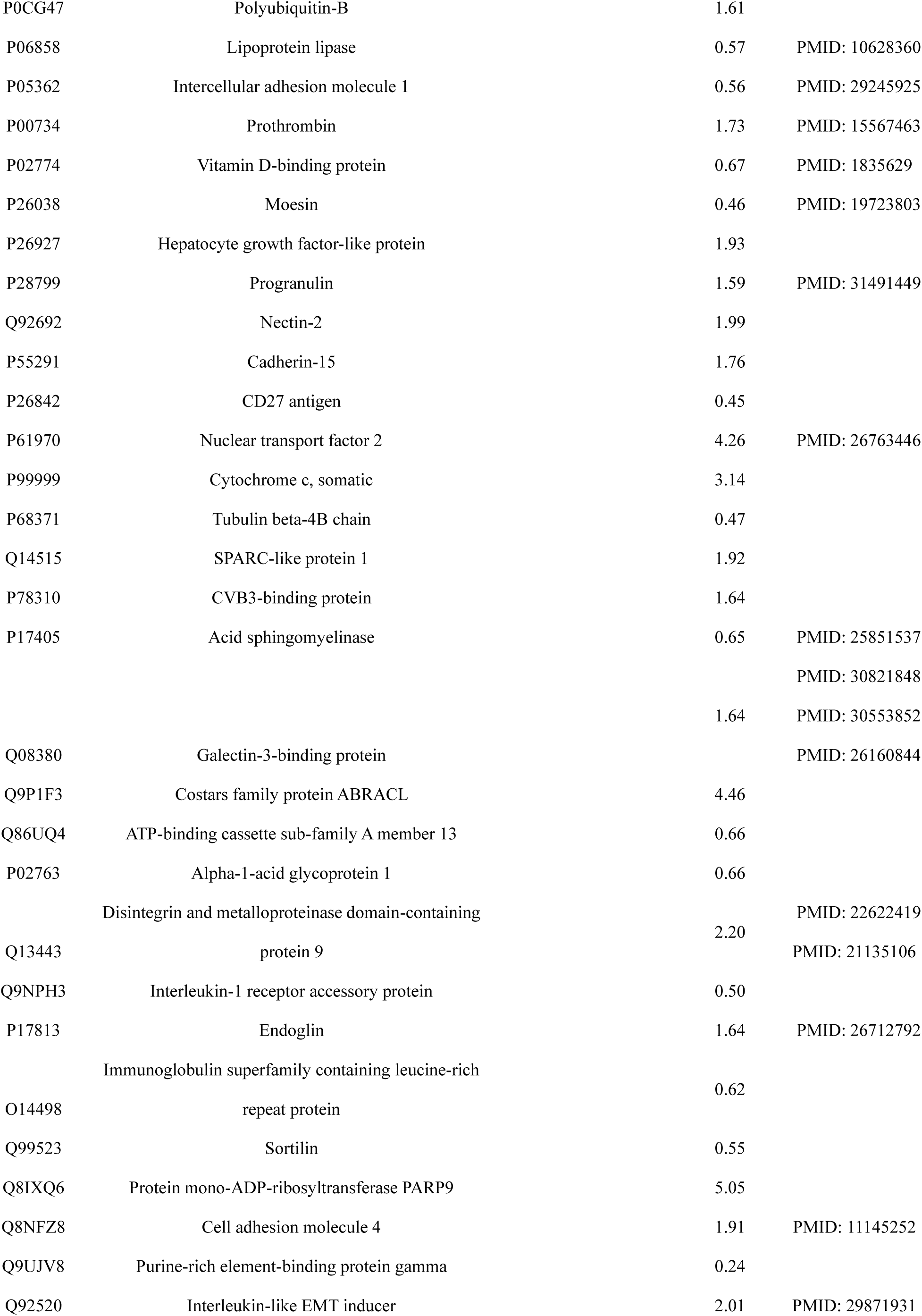
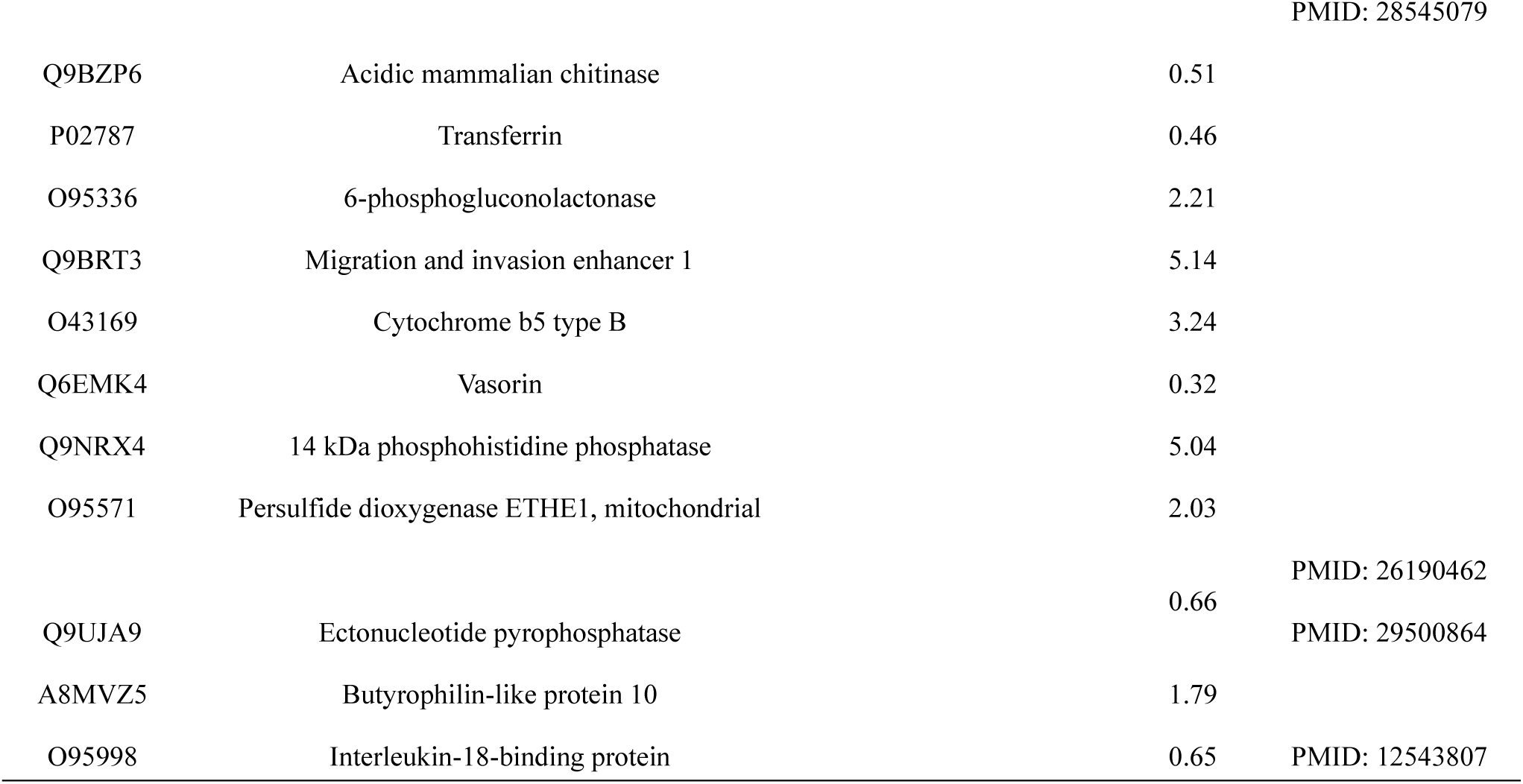
Differential proteins in the B16 tumor-bearing mice (p<0.05)

**Table 6B.**
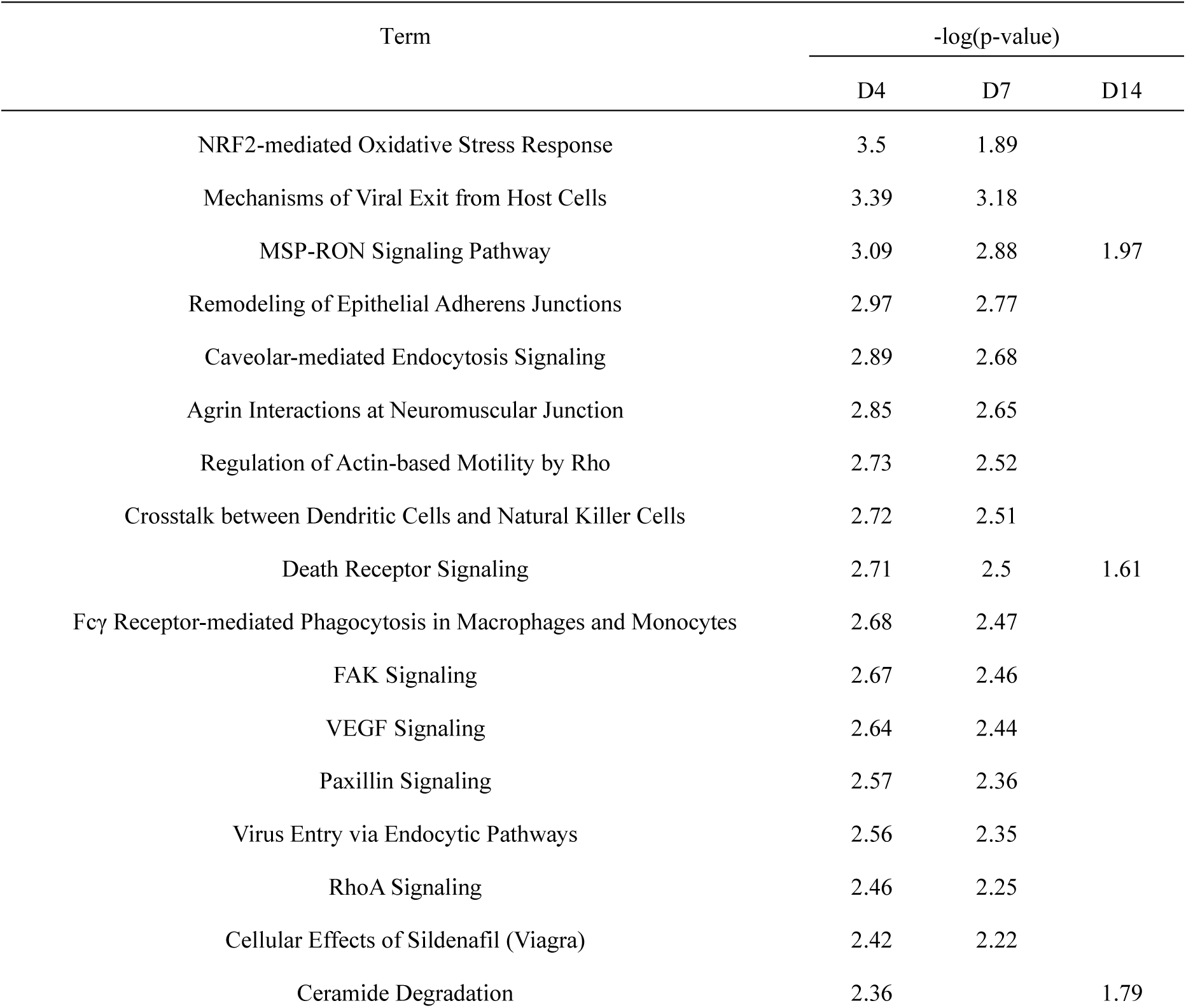
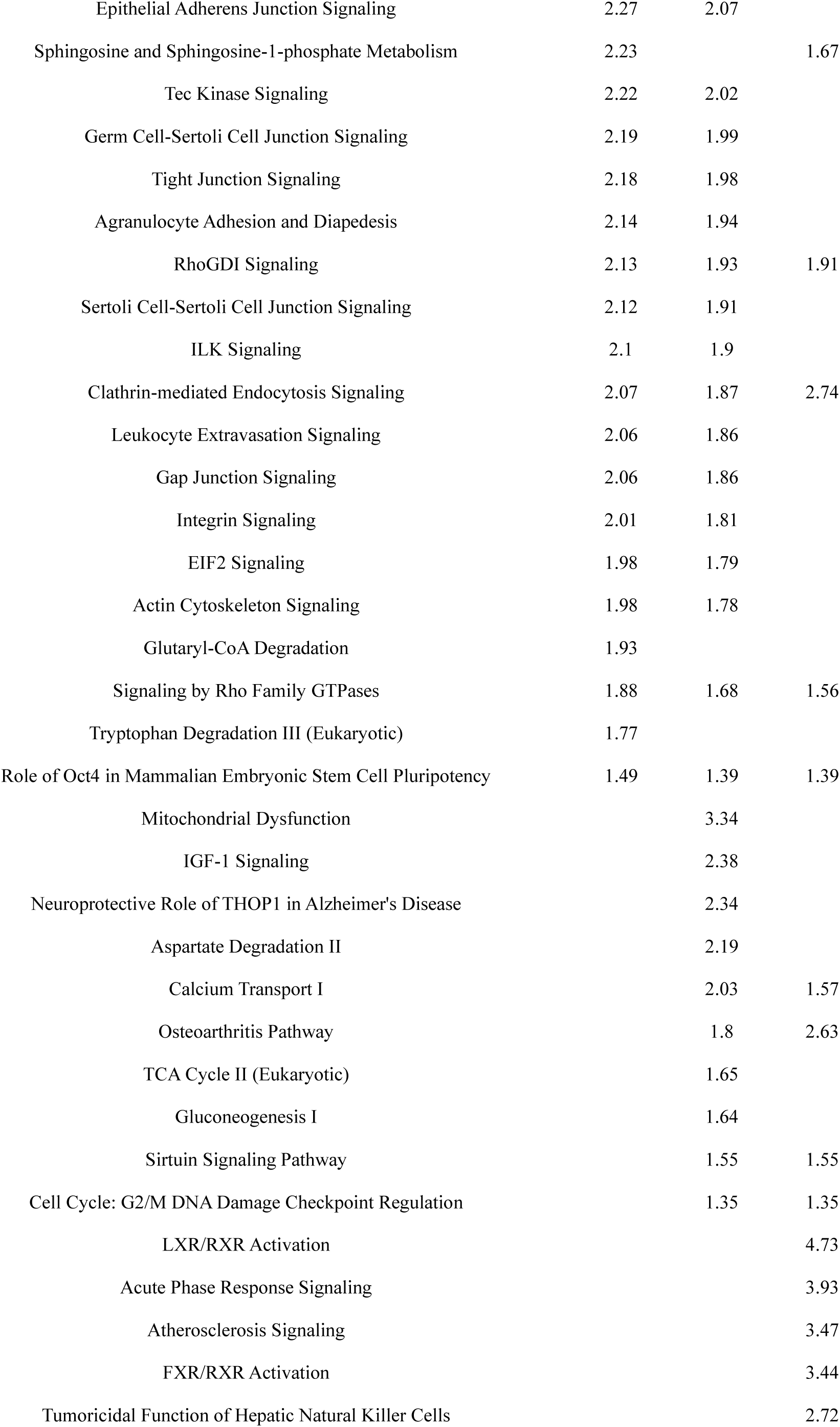
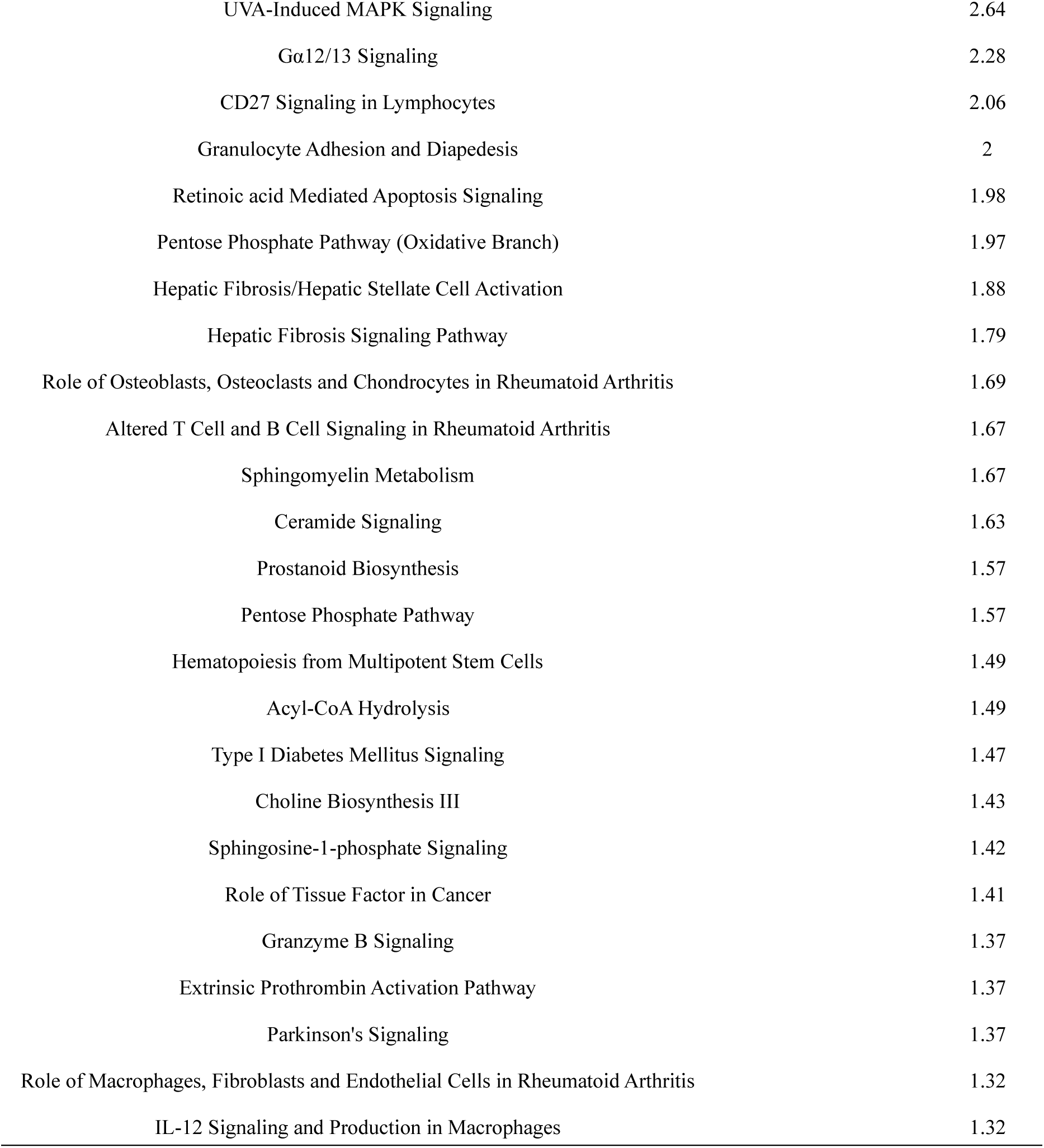
Pathway analysis of differential proteins in the B16 tumor-bearing mice (p<0.05)

#### 2.3.2 RM-1 prostate cancer-bearing mice

Analysis of biological processes showed metabolic and inflammatory processes on Day 7 and Day 22 and bone development on Day 15, suggesting that this process may be related to tumor bone metastasis and is consistent with the trend of bone metastasis in prostate cancer (Figure 10A). Analysis of cell composition also revealed that the differential proteins were secreted proteins, mostly from the extracellular matrix and the cell surface (Figure 10B). Analysis of the molecular function revealed the activity of the channel protein (Figure 10C). Similarly, the pathway analysis showed that the pathways at different time points had their own characteristics, and some of them were reported to be associated with prostate cancer (Table 7). For example, before the tumor was palpable, FXR/RXR activation, LXR/RXR activation[83], mitochondrial dysfunction, acute phase response signaling and other pathways were significantly enriched. It has been reported that the activation of the LXR/RXR and FXR/RXR pathways may alleviate inflammation. RXR is a nuclear hormone receptor of the retinoid A receptor family, which is usually used with LXR and FXR receptors. These receptors usually exist in biological and pathological pathways related to glucose, lipid homeostasis and the inflammatory response[83]. The proteomic study of urine in prostate cancer patients with a Gleason score of 6.9 ± 1.1 showed that LXR/RXR is activated and the signal of the acute-phase response is significantly downregulated[84]. Clathrin-mediated endocytosis is associated with the migration of prostate cancer cells[85]. The regulation of Rho on actin movement is significantly enriched after the tumor is palpable. It has been reported that the RhoC gene can affect cell migration by regulating the actin cytoskeleton, thus affecting the invasion and metastasis of malignant tumors[86].

**Figure 10A.**
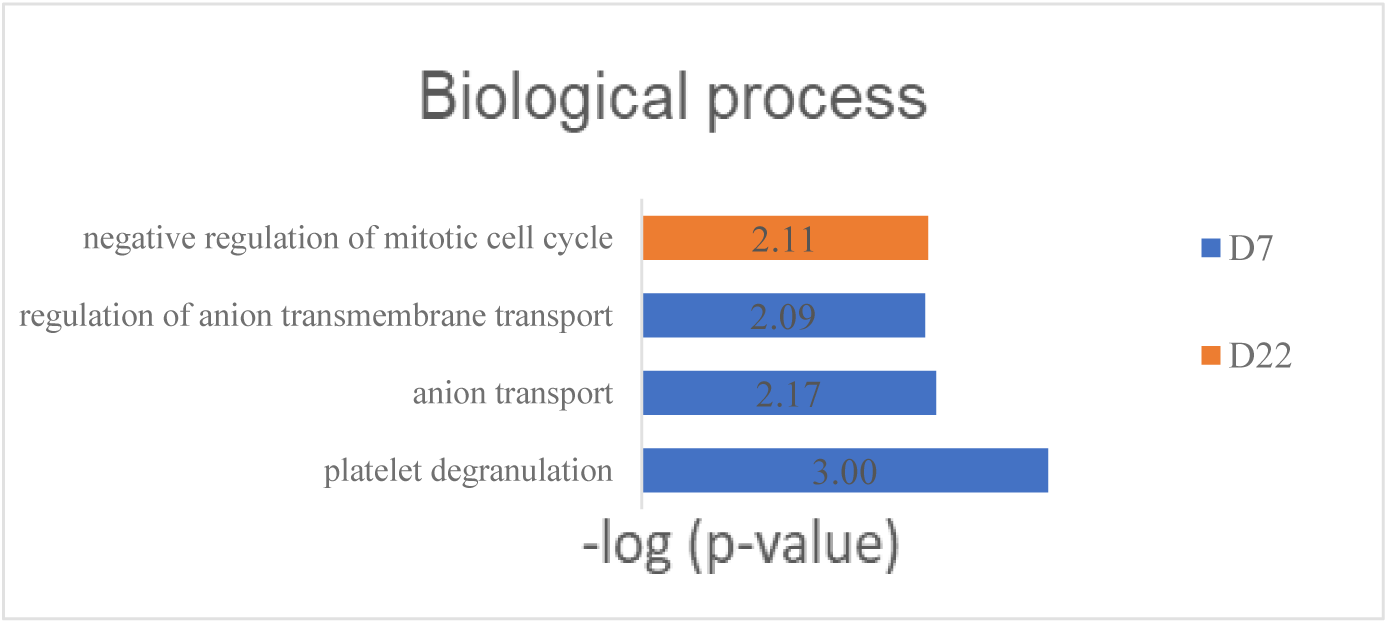
Biological process analysis of differential proteins in the RM-1 tumor-bearing mice

**Figure 10B.**
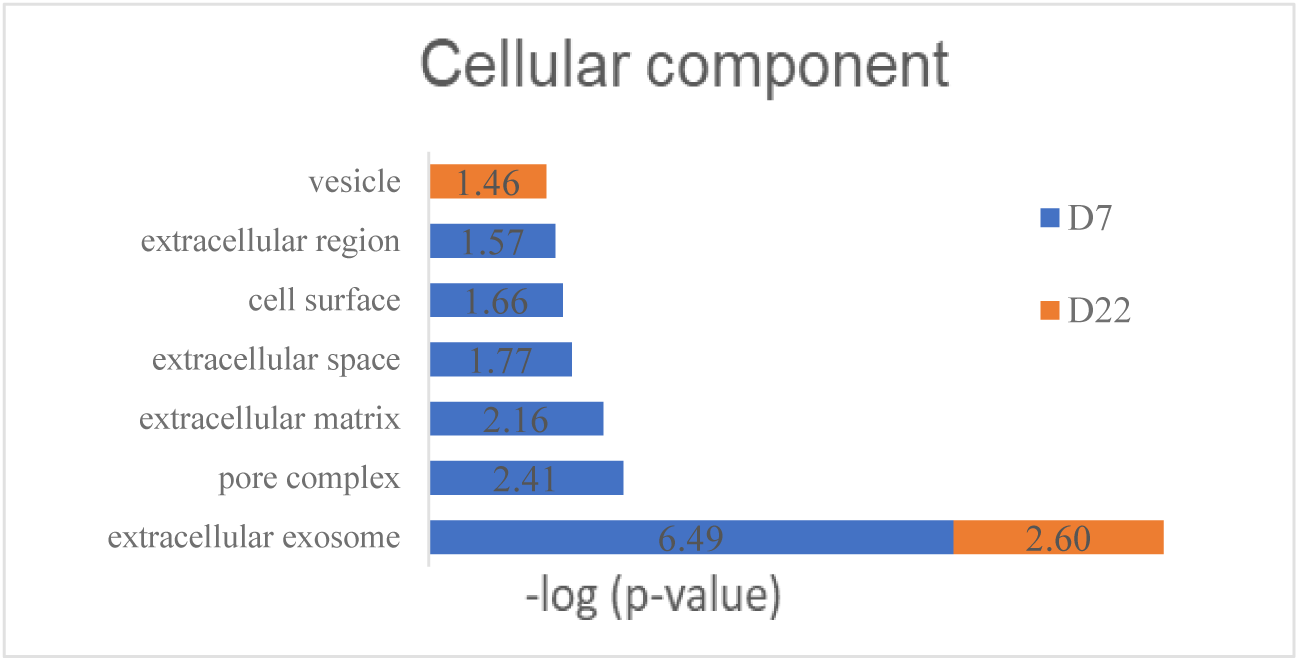
Cell component analysis of differential proteins in the RM-1 tumor-bearing mice

**Figure 10C.**
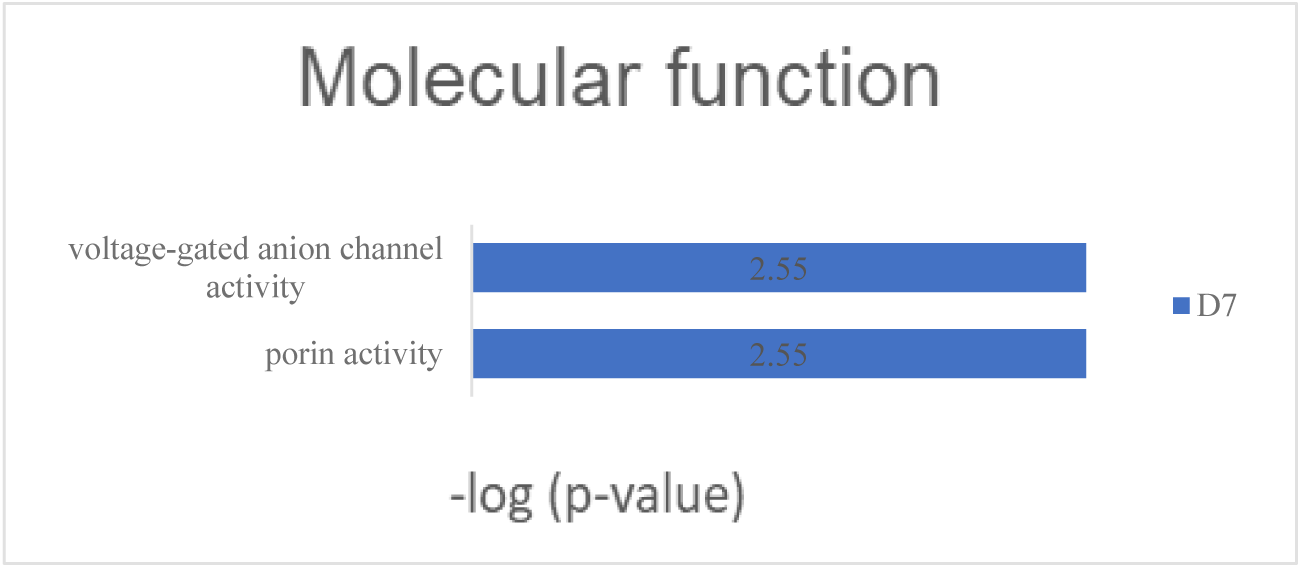
Biological process analysis of differential proteins in the RM-1 tumor-bearing mice

**Table 7.**
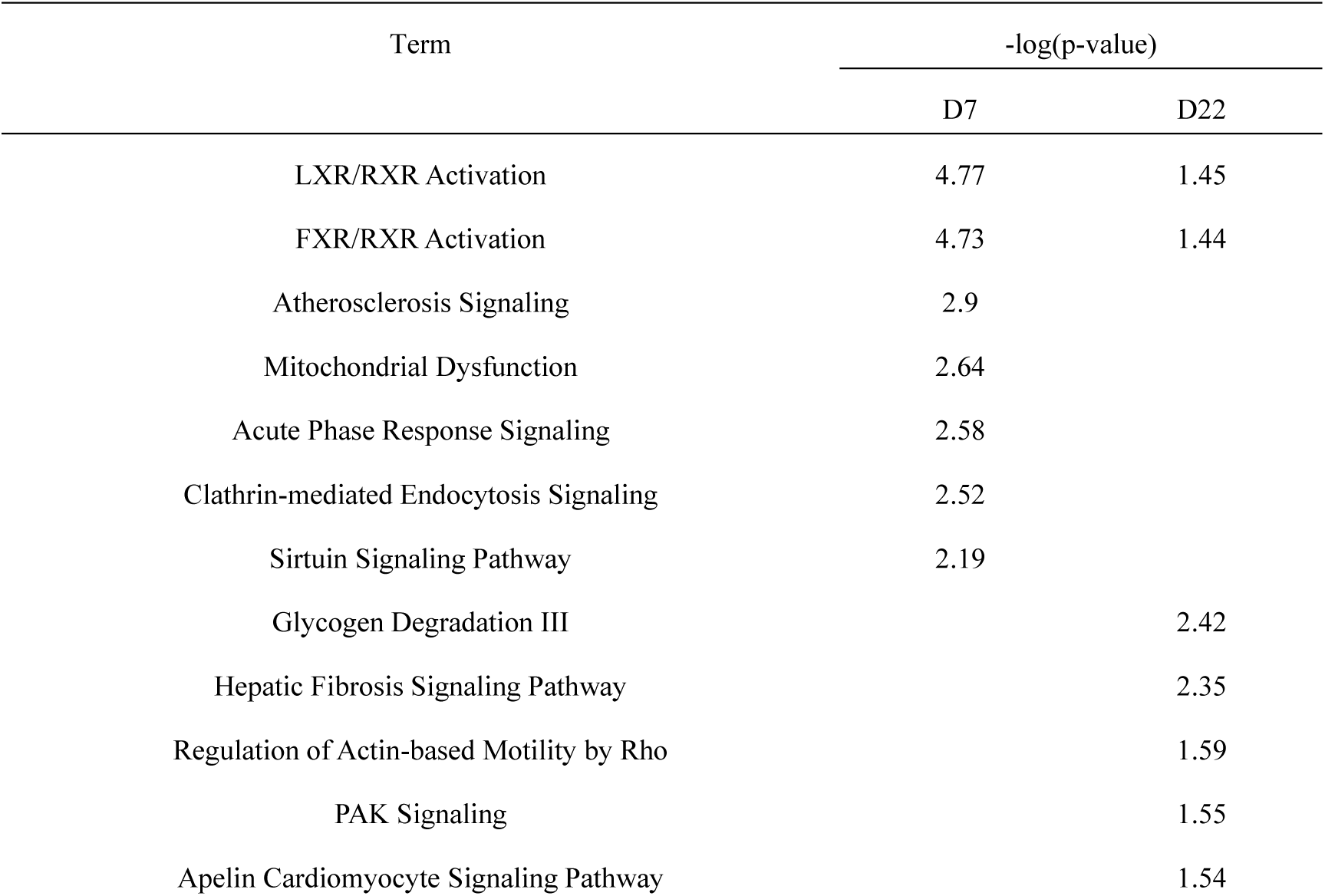
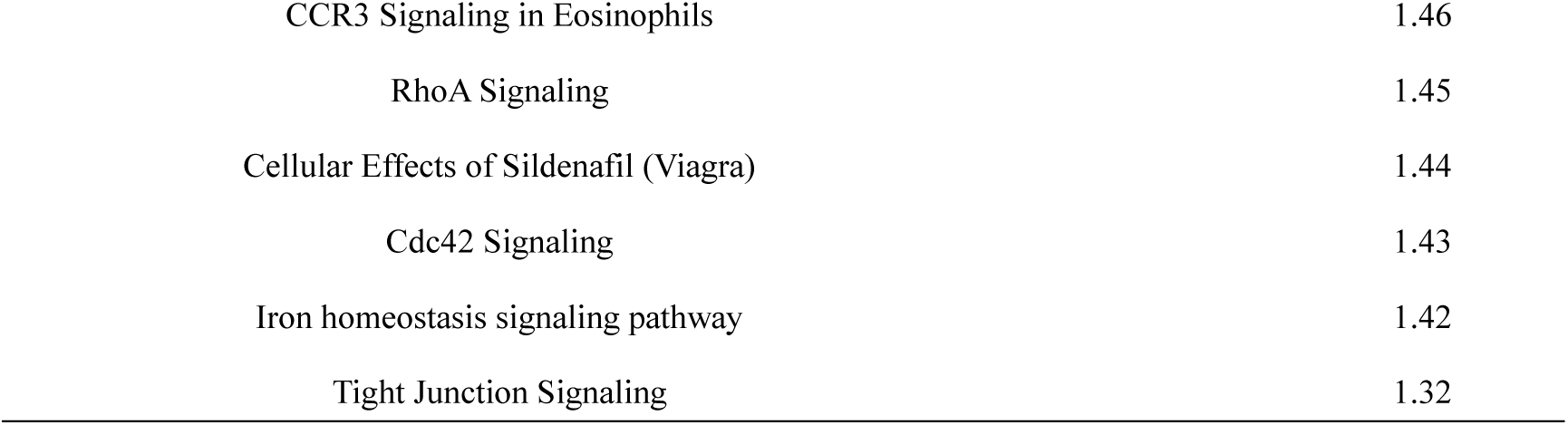
Pathway analysis of differential proteins in the RM-1 tumor-bearing mice

A total of 42 homologous differential proteins were identified with the condition of “fold change > 1.5, p value < 0.05”, of which 23 were reported to be associated with prostate cancer (Table 8, Figure 11).

**Table 8A.**
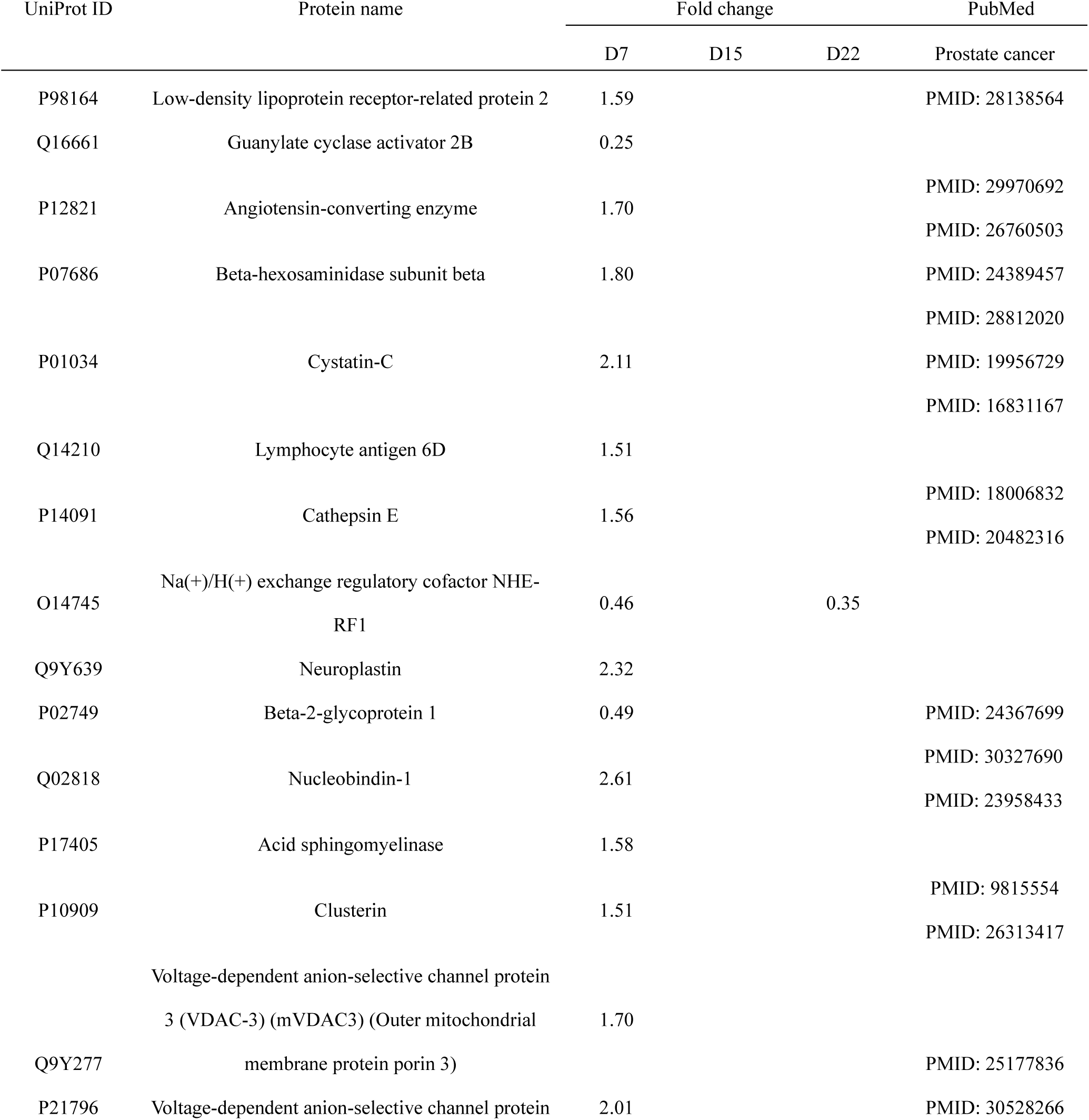
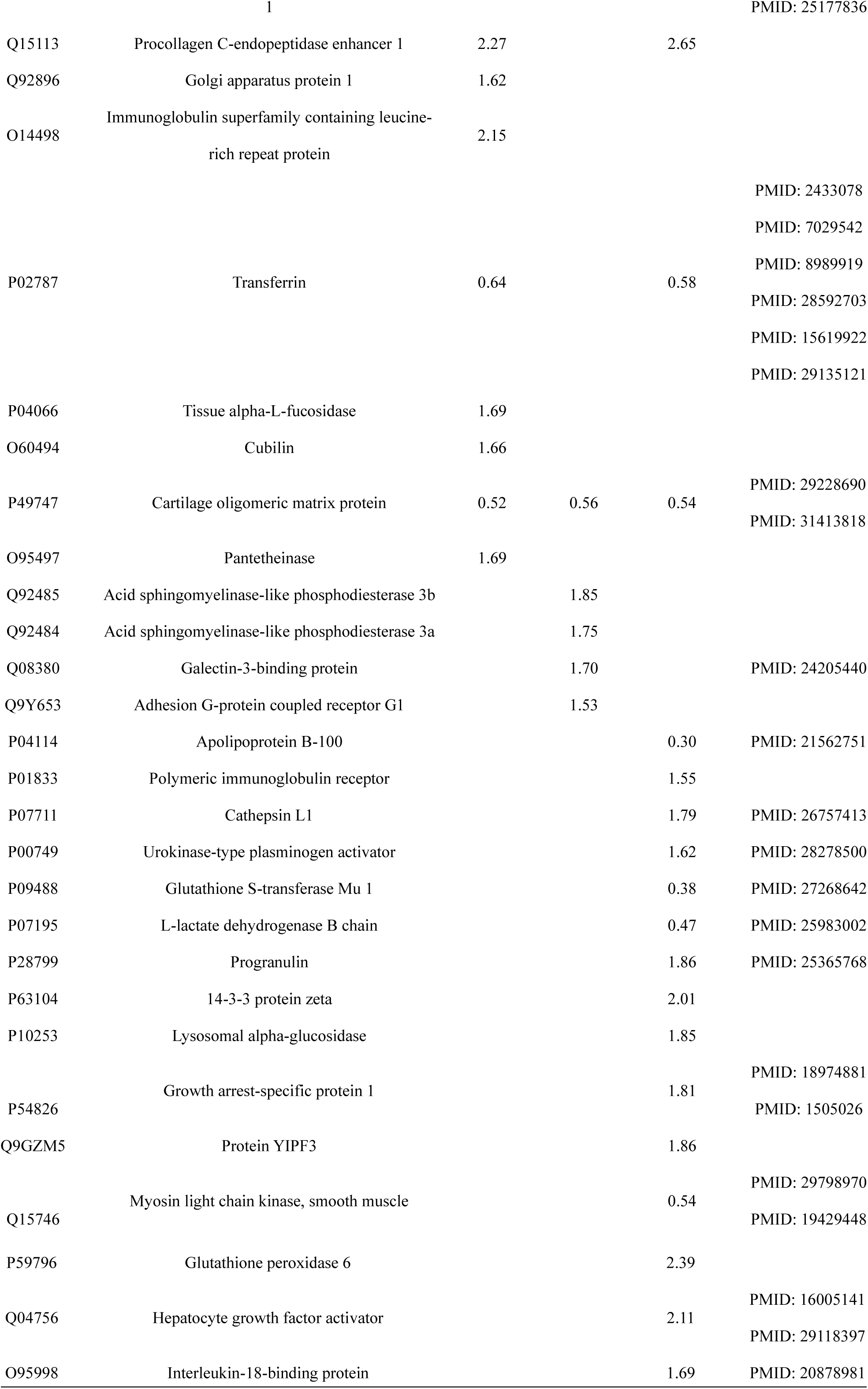
Differential proteins in the RM-1 tumor-bearing mice (p<0.05)

**Table 8B.**
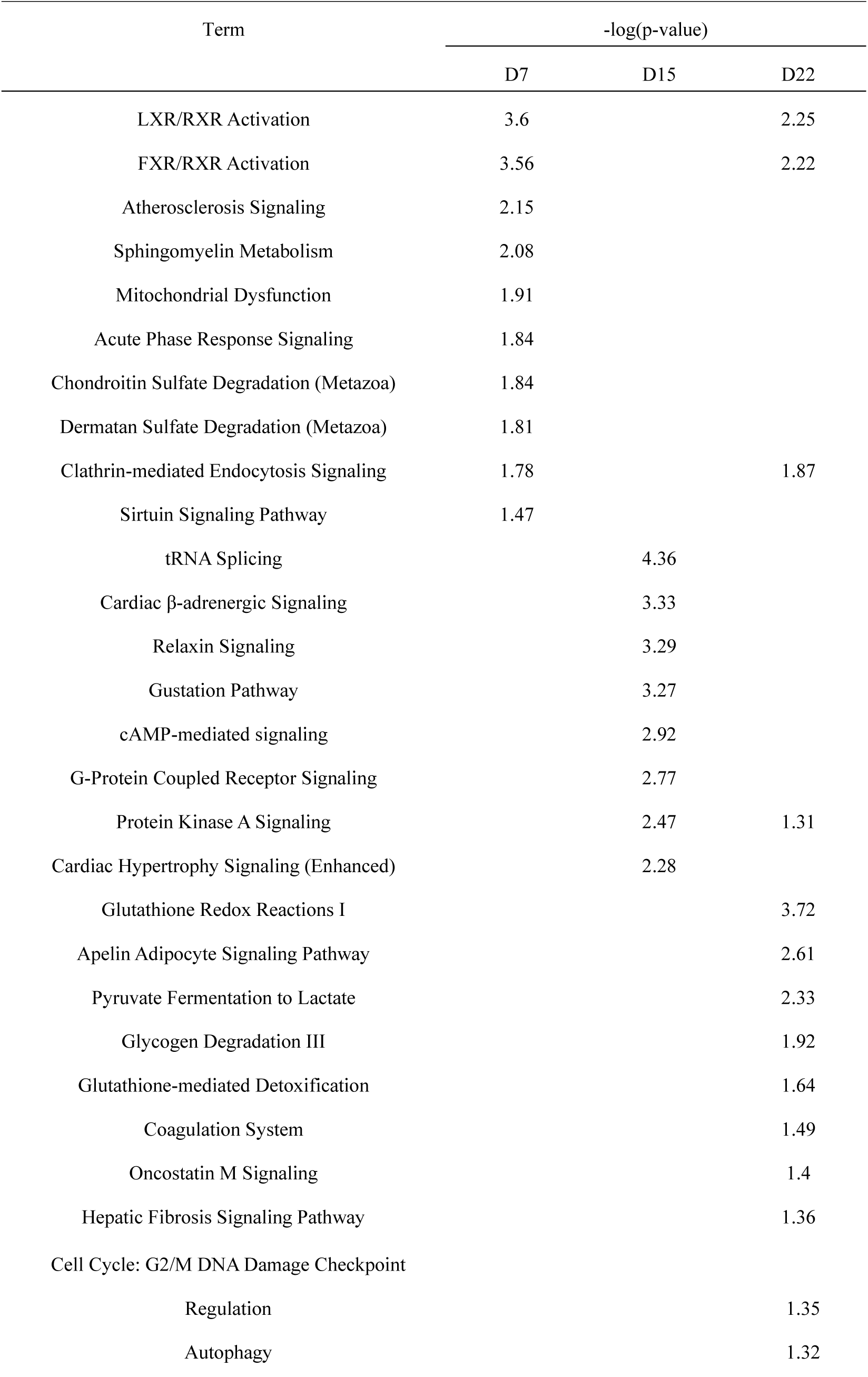
Pathway analysis of differential proteins in the RM-1 tumor-bearing mice (p<0.05)

**Figure 11A.**
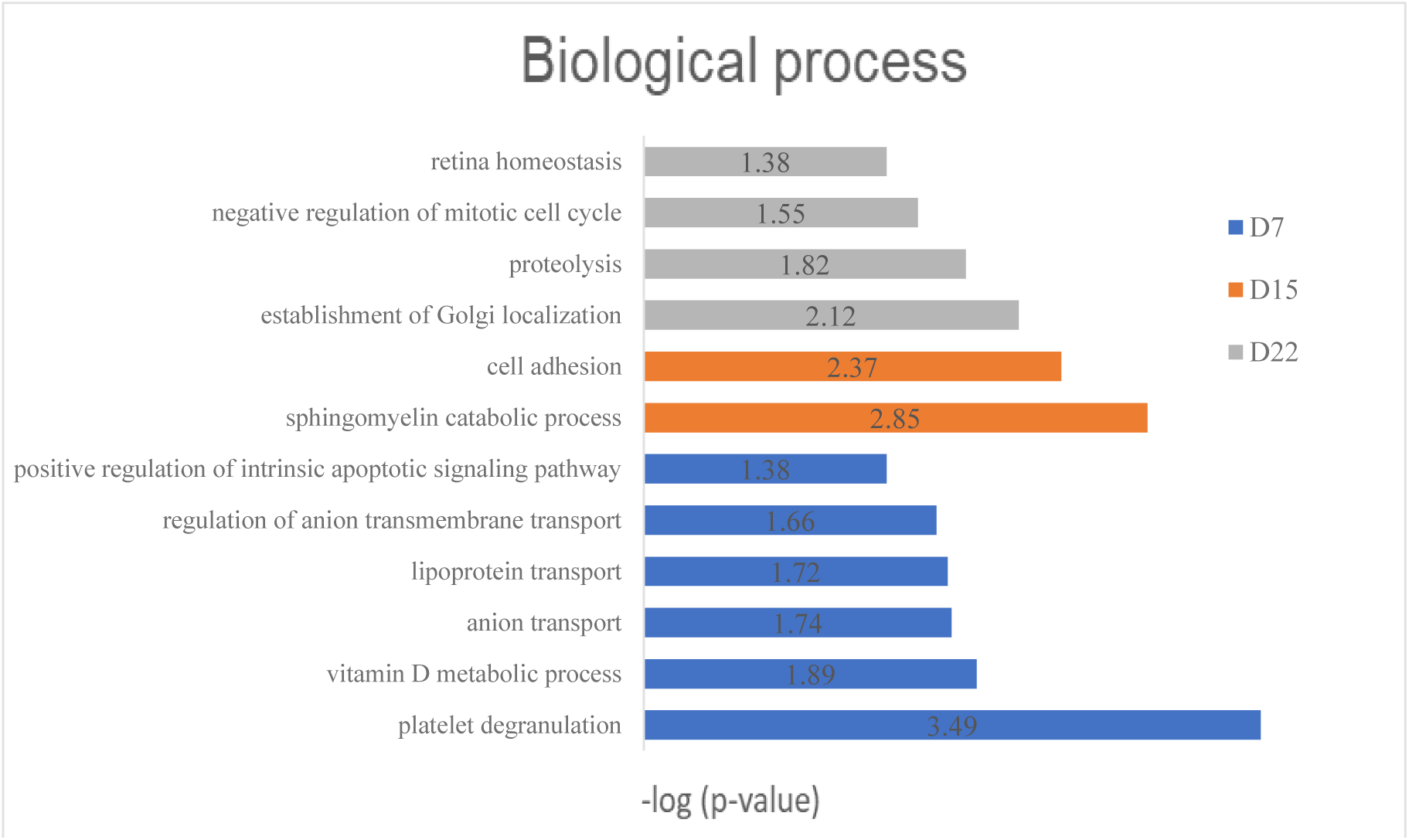
Biological process analysis of differential proteins in the RM-1 tumor-bearing mice (p<0.05)

**Figure 11B.**
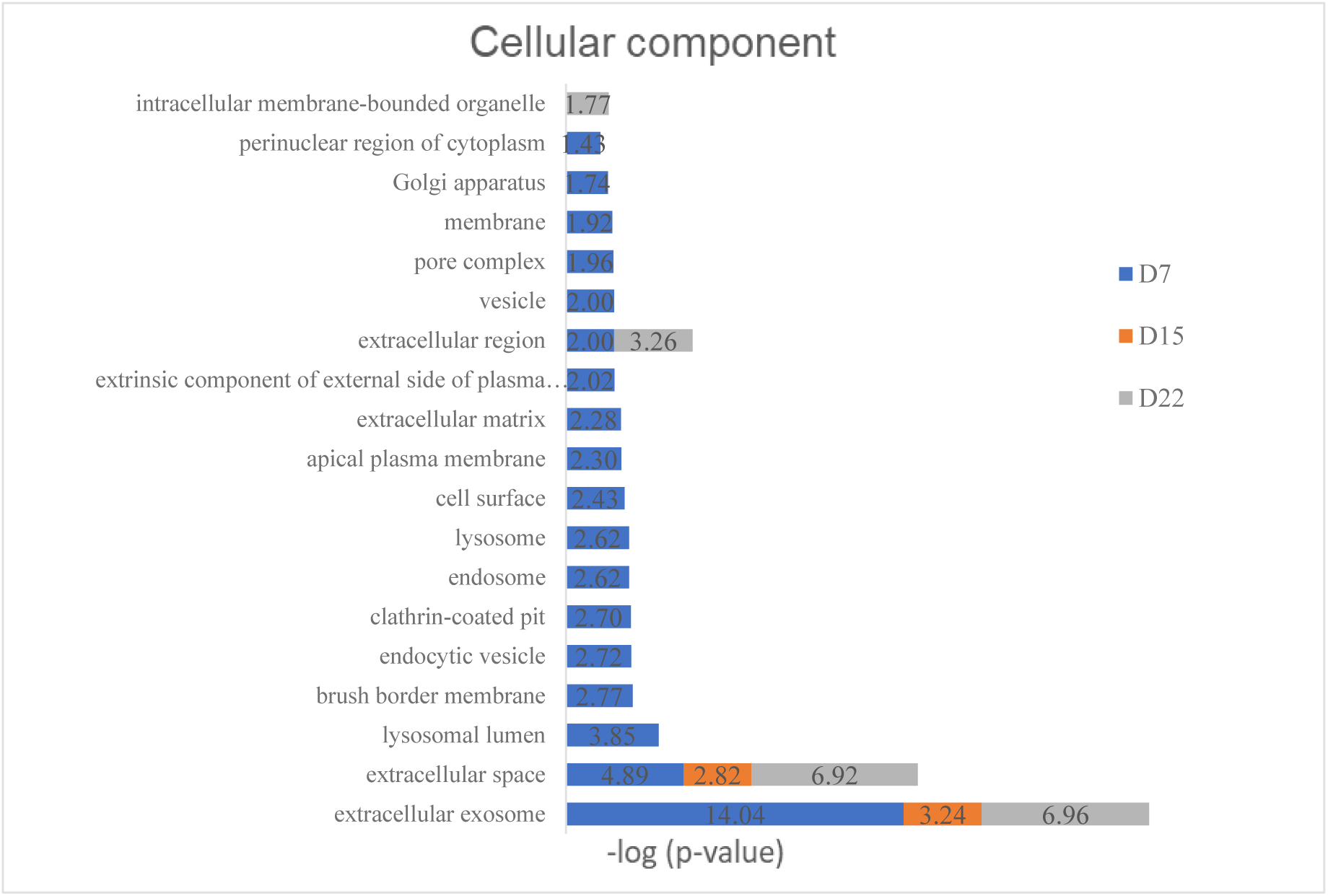
Cell component analysis of differential proteins in the RM-1 tumor-bearing mice (p<0.05)

**Figure 11C.**
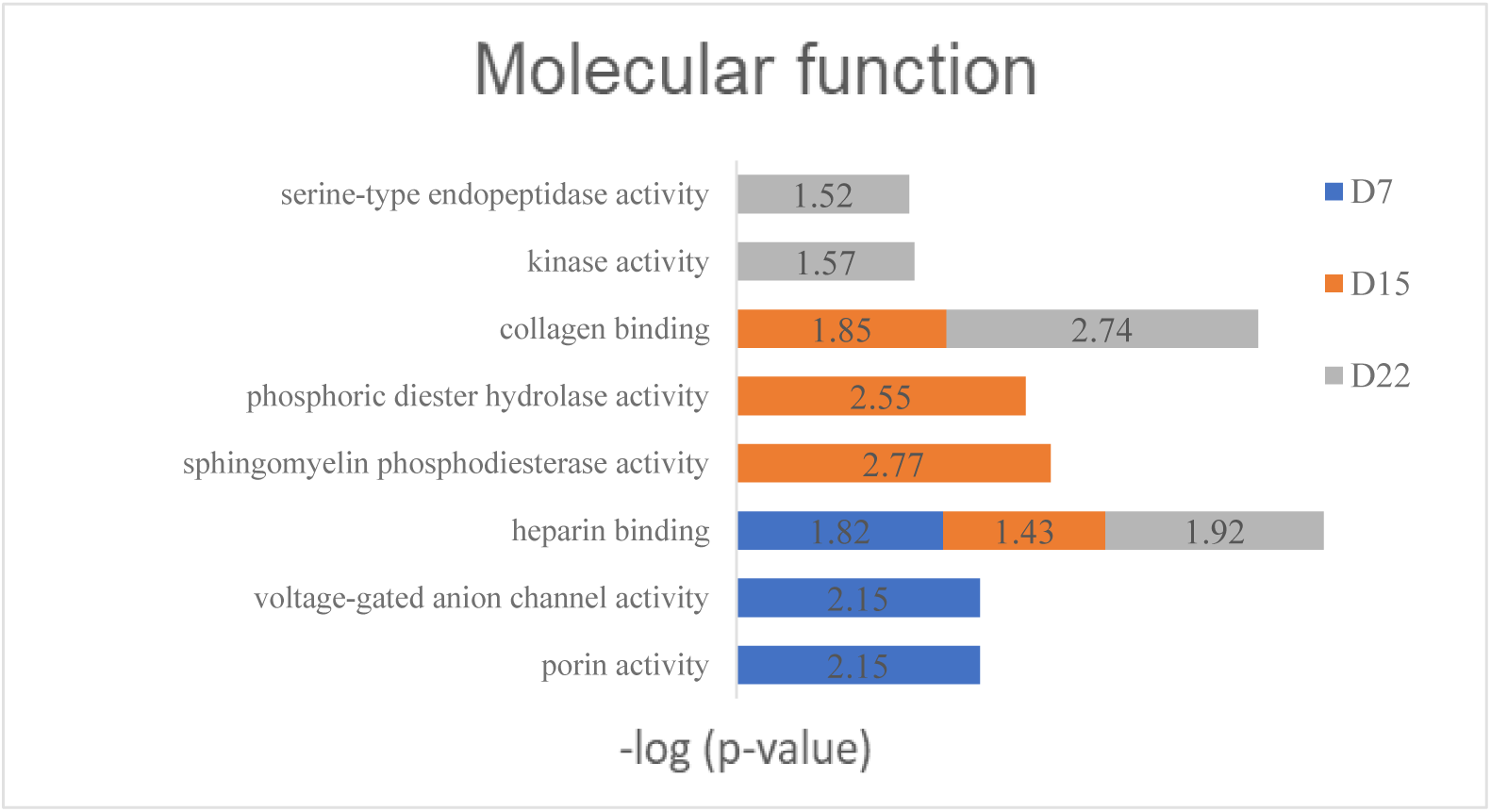
Molecular function analysis of differential proteins in the RM-1 tumor-bearing mice (p<0.05)

## 3 Conclusion

Random group analysis of samples can independently select better screening conditions according to the specific situation of experimental data, which is conducive to finding a group of differential proteins with higher reliability, and combination or linkage of differential proteins can further improve the reliability of markers. Furthermore, the reliability of differential proteins is related to the development of disease and the number of animals. To maximize the reliability of urine proteome changes, researchers can increase the number of experimental animals in future studies of urine biomarkers, particularly for early studies of diseases or diseases with little variation.

The results of this study showed that the urine proteome could reflect changes in the development of melanoma and prostate cancer cells. Before the tumor was detectable, significantly differential proteins were identified in the urine of both the B16 melanoma and RM-1 prostate tumor-bearing models. Moreover, there are major differences among the tumor-bearing models, suggesting that urine has good potential for early diagnosis, differentiation of different tumor types and development. Eventually, the different combinations of biomarkers can provide new ideas and clues for the early diagnosis of a tumor, which has the value in large-scale clinical research.

